# Control of Mast Cell Regulated Exocytosis by Munc13 Proteins

**DOI:** 10.1101/202473

**Authors:** Elsa M. Rodarte, Marco A. Ramos, Alfredo J. Davalos, Daniel C. Moreira, David S. Moreno, Eduardo I. Cardenas, Alejandro I. Rodarte, Youlia Petrova, Sofia Molina, Luis E. Rendon, Elizabeth Sanchez, Keegan Breaux, Alejandro Tortoriello, John Manllo, Erika A. Gonzalez, Michael J. Tuvim, Burton F. Dickey, Alan R. Burns, Ruth Heidelberger, Roberto Adachi

## Abstract

Mast cells (MCs) are involved in pathogen defense and inflammatory reactions. Upon stimulation, they release substances stored in their granules via regulated exocytosis. In other cell types, Munc13 proteins play essential roles in regulated exocytosis. We found that MCs express Munc13-2 and -4, and we studied their roles using global and conditional knockout (KO) mice. In a model of systemic anaphylaxis, we found no difference between WT and Munc13-2 KO mice, but global and MC-specific Munc13-4 KO mice developed less hypothermia. This protection correlated with lower plasma histamine levels and histological evidence of defective MC degranulation, and not with changes in MC development, distribution, numbers or morphology. In vitro assays revealed that the defective MC response in the absence of Munc13-4 was limited to regulated exocytosis, leaving other MC secretory effector responses intact. Single cell capacitance measurements in MCs from mouse mutants with different expression levels of Munc13-4 in their MCs showed that as levels of Munc13-4 decrease, the rate of exocytosis declines first, and the total amount of exocytosis follows. A requirement for Munc13-2 in MC exocytosis was revealed only in the absence of Munc13-4. Electrophysiology and electron microscopy studies showed that the number of multigranular compound events (granule-to-granule homotypic fusion) was severely reduced in the absence of Munc13-4. We conclude that while Munc13-2 plays a minor role, Munc13-4 is essential for regulated exocytosis in MCs, and that this MC effector response is required for a full IgE-mediated anaphylactic response.

---

During exocytosis, the membrane of a secretory vesicle fuses with the plasma membrane, allowing the release of vesicular contents into the extracellular space and the incorporation of vesicle membrane components into the plasma membrane (1). Exocytosis can be constitutive or regulated (2). In constitutive exocytosis, newly formed products are secreted as they are synthesized, and the amount of secreted product is controlled by the rate of expression of the vesicular cargo. In contrast, in regulated exocytosis, the formed products are stored in secretory vesicles (e.g., MC granules) and released upon stimulation using diacylglycerol (DAG) and Ca^2+^ as second messengers (3,4). The amount of secreted product is controlled by the rate and number of vesicle to plasma membrane fusion events. Regulated exocytosis can adopt various forms. In single-vesicle exocytosis, individual secretory granules fuse with the plasma membrane. In sequential compound exocytosis, these primary fused vesicles become targets for secondary fusion events with vesicles lying deeper in the cell. In multigranular compound exocytosis, secretory vesicles fuse homotypically with each other inside the cell before fusing heterotypically with the plasma membrane (5). Some cells (e.g., MCs) use all three forms of regulated exocytosis (6).

Regulated exocytosis involves the generation of secretory vesicles and their transport towards the plasma membrane. Then, tethering and docking establish physical proximity between the vesicle and plasma membrane. The final event involves the fusion of both membranes (1), which requires the assembly of complexes between the SNARE (<u>s</u>oluble <u>N</u>-ethylmaleimide-sensitive factor <u>a</u>ctivated protein <u>r</u>eceptor) domains of proteins on the vesicular (vesicle associated membrane protein [VAMP]) and target membranes (syntaxin [Stx] and synaptosomal-associated protein 25 [SNAP25]) (7), and is highly regulated by complexin and synaptotagmin (8). The physical proximity established by docking is not sufficient to drive fusion, additional biochemical events are required to render the vesicles competent for Ca^2+^-triggered fusion in an intermediate step known as priming (9,10).

The mammalian homologs of *C. elegans* uncoordinated gene 13 (Munc13) are essential for docking (11) and priming (12,13). All isoforms contain a MUN domain, which in Munc13-1 is required for making the SNARE domain of Stx available to interact with those of VAMP and SNAP25 (14,15) in the correct configuration (16). Munc13 proteins also contain two or three C2 domains. The C2A domain regulates the function of Munc13 through homodimerization or binding to RIM (Rab3-interactin molecule) (17,18). C2B and C2C help to bridge the vesicular and plasma membranes (18,19). Some Munc13 isoforms are also have a C1 domain (18,20), and binding of DAG to C1 and of Ca^2+^ to C2B regulate the final assembly of the SNARE complex (21). Neuronal loss of Munc13-1 severely impairs neurotransmitter release and drastically reduces the number of fusion-ready vesicles (22). Munc13-4 is ubiquitously expressed, and most studies have focused on lymphocytes because its absence in humans causes familial lymphohistiocytosis type 3 (23). Compared to Munc13-1 and -2, Munc13-4 lacks the C2A and C1 domains (24). Munc13-4 binds in vitro to the SNARE domains of Stx-1, 4 and 11 (25), and facilitates the fusion of Rab7^+^ secretory granules with Rab11^+^ endosomes in RBL-2H3 cells (26).

MCs can be activated by allergens, complement, cytokines, growth factors, venoms, and other secretagogues (27). One MC effector response is degranulation, in which mediators stored in their large metachromatic granules are released via regulated exocytosis (28,29). Stimulation also activates the transcription of multiple cytokines, chemokines and growth factors, which are synthesized in the ER and then exported via constitutive exocytosis from the Golgi to the plasma membrane. In addition, the rise in intracellular Ca^2+^ induces the enzymatic processing of arachidonic acid into eicosanoids, mainly PGD_2_ and LTC_4_, which are exported through membrane transporters (30,31).

We hypothesized that the robust degranulation kinetics of MCs would allow us to test with high resolution the requirements of Munc13 proteins in non-neuronal cell exocytosis. To this end, we used single-cell, cell population and whole-animal assays. We found that Munc13-4 regulates the amount and rate of exocytic events, that it is specifically required for MC regulated exocytosis but not for other MC effector responses, that it mediates homotypic fusion in multigranular compound exocytosis, and that animals with a selective deficiency of Munc13-4 in their MCs could not mount a full anaphylactic response.

## RESULTS

### Expression of Munc13 proteins in MCs and generation of Munc13-4-deficient mice

We found that C57BL/6J (B6) mature peritoneal MCs express Munc13-2 and -4 (Fig. 1*A*), and decided to target both isoforms. We obtained Munc13-2 global KO mice (32) and generated Munc13-4 global and conditional KO lines (Fig. 1*B*). We flanked the exon containing the transcription initiation codon of *Unc13d* with two loxP sequences (“floxed” or F allele) and removed it in MCs (Δ allele) using mice that express Cre recombinase under the control of the *Cma1* (chymase 1) locus (33). We also crossed Munc13-4^F/F^ with CMV-Cre mice (34) to obtain germline deletion (– allele). PCR of genomic DNA with primers surrounding exon 3 confirmed our genetic manipulations (Fig. 1 *C*). The resulting mutant mice (Munc13-4 +/−, −/−. F/F and Δ/Δ) were all viable and fertile, transmission of the mutant alleles followed a Mendelian pattern, and we found no gross anatomical abnormalities or survival differences when raised in a pathogen-free facility.

**Figure 1.**
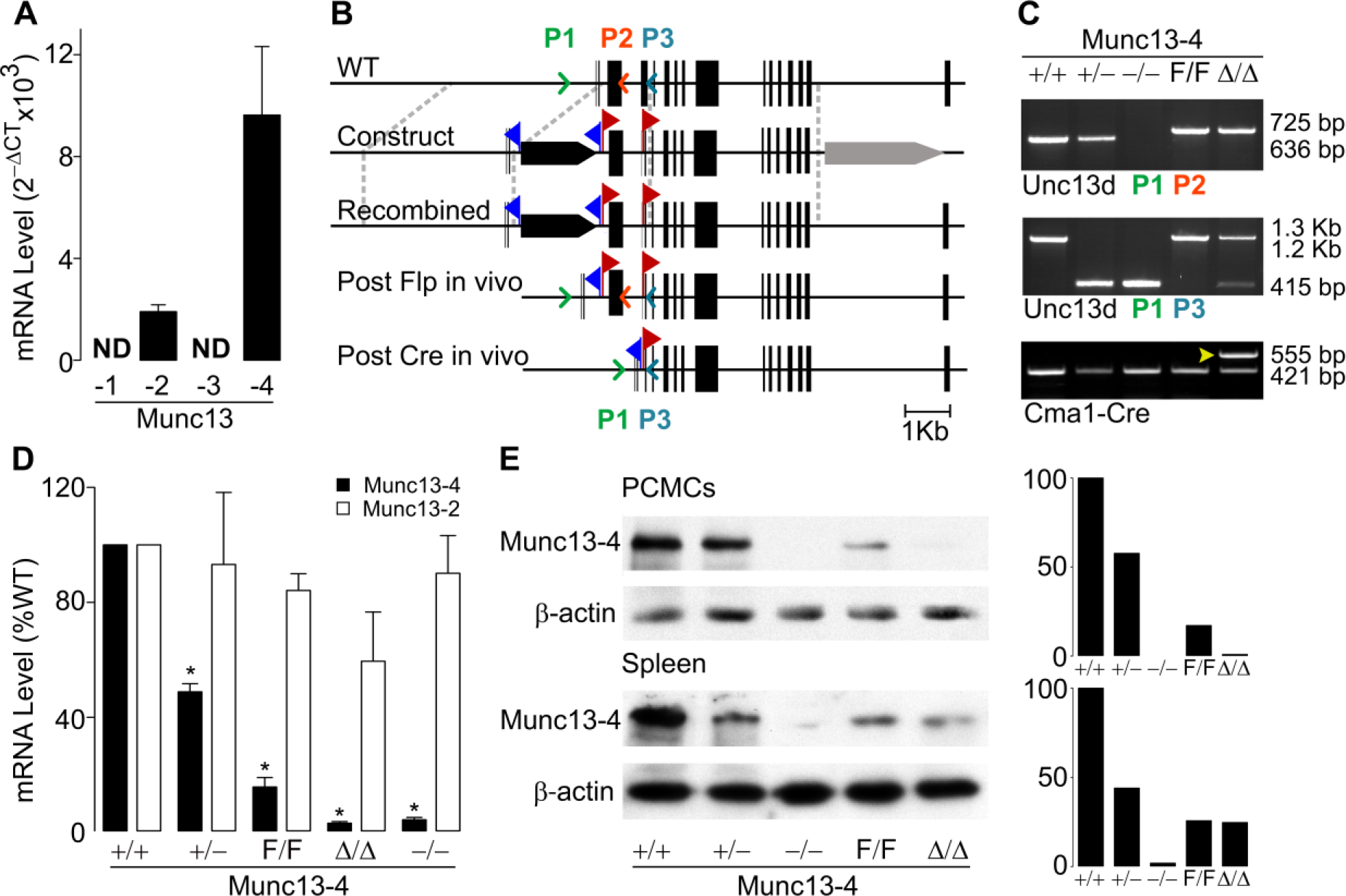
Expression of Munc13 proteins in MCs from WT and mutant mice. *A*, qPCR of B6 peritoneal MCs for all Munc13 paralogs relative to β-actin. ND, not detected; N = 4. *B*, generation of the Munc13-4 global and conditional KO mice. Black bars, exons; blue flags, location of FRT sites; red flags, location of loxP sites; black arrow, PGK-Neo; grey arrow, PGK-TK; color arrowheads, location of primers for genotyping. *C*, PCR of genomic tail DNA amplified with the primers in panel B. Arrow head, Cre^+^ lane. *D*, qPCR for Munc13-2 and Munc13-4 of peritoneal MCs from different Munc13-4 mutant mice. Values are normalized to β-actin and then to levels in Munc13-4^+/+^ MCs. Bar, mean; error bar, SEM. N = 4; * = p ≤ 0.001 compared to Munc13-4^+/+^. *E*, representative immunoblot of PCMC and spleen lysates for Munc13-4. β-actin was used as loading control. Right insets, densitometry relative to β-actin and Munc13-4^+/+^.

The efficacy of Cre recombination in peritoneal MCs using a surrogate locus was > 99% (supplemental Fig. S1), so we used peritoneal MCs in most of our studies. Quantification of transcripts from FACS-sorted peritoneal MCs demonstrated the predicted absence of Munc13-4 transcripts in MCs from Munc13-4^−/−^ and Munc13-4^Δ/Δ^ mice, and ~50% reduction in those from Munc13-4^+/−^ mice. Also, there was no compensatory overexpression of Munc13-2 in the absence of Munc13-4 (Fig. 1*D*). The reduction of Munc13-4 transcript levels in Munc13-4^F/F^ MCs to ~15% was unexpected. By 3′ rapid amplification of cDNA ends (RACE), we confirmed that the transcript present in Munc13-4^F/F^ mice was a WT Munc13-4 transcript. Immunoblots of lysates from PCMCs (peritoneal cell-derived MCs) and different organs (only spleen is shown) confirmed the reduced expression in Munc13-4^F/F^ mice and the specificity of Cre-mediated deletion: Munc13-4^Δ/Δ^ animals had the same levels of Munc13-4 as Munc13-4^F/F^ mice in all tissues except in MCs, where it was absent (Fig. 1*E*). Because Munc13-4^FF^ mice were hypomorphic, Munc13-4^Δ/Δ^ mice were strictly compared to Munc13-4^F/F^ littermates in all experiments.

### Munc13-4 is required for a full anaphylactic response

In small mammals, the massive peripheral vasodilation characteristic of anaphylaxis causes rapid loss of body heat, so core body temperature accurately reflects the severity of the anaphylactic response. Hypothermia in this model of IgE-dependent anaphylaxis depends on histamine (35). Given that histamine is released by MCs via regulated exocytosis, we tested if the global or MC-specific deletion of Munc13-4 affected this allergic response. We sensitized mice with IgE anti-DNP (2,4-dinitrophenol), challenged them with DNP-human serum albumin (DNP-HSA) and monitored their core body temperature (Fig. 2*A*). As negative controls we introduced Munc13-4^+/+^ mice that were challenged with a sham solution. To control for anaphylactoid reactions to DNP-HSA we used non-sensitized but challenged B6 mice and no reactions were observed (data not shown). The lack of response in MC-deficient Kit^W-sh/W-sh^ mice (W^sh^) confirmed the MC-dependency of this model. The absence of Munc 13-2 did not affect the hypothermic response. However, there was significantly less drop in temperature in Munc13-4^−/−^ mice compared to Munc13-4^+/+^ littermates, suggesting that a component of this reaction depends on Munc 13-4. We recorded a similar difference between Munc13-4^Δ/Δ^ and Munc13-4^F/F^ mice, indicating that this Munc13-4-dependent component is limited to MCs (Fig. 2*C*). The partial deficit of Munc13-4 expression in Munc13-4^+/−^ and Munc13-4^F/F^ mice did not modify the nadir or recovery phase of the anaphylactic response, but it affected the early phase (Fig. 2*B*). Based on these findings we decided to drop the Munc13-2^−/−^ mice from most of our studies and continue studying our Munc13-4 mutant mice. Also, because we did not find a difference between Munc13-4^−/−^ and Munc13-4^Δ/Δ^ mice in this and most other experiments, we opted not to compare Munc13-4^+/−^ with Munc13-4^+/Δ^ mice.

**Figure 2.**
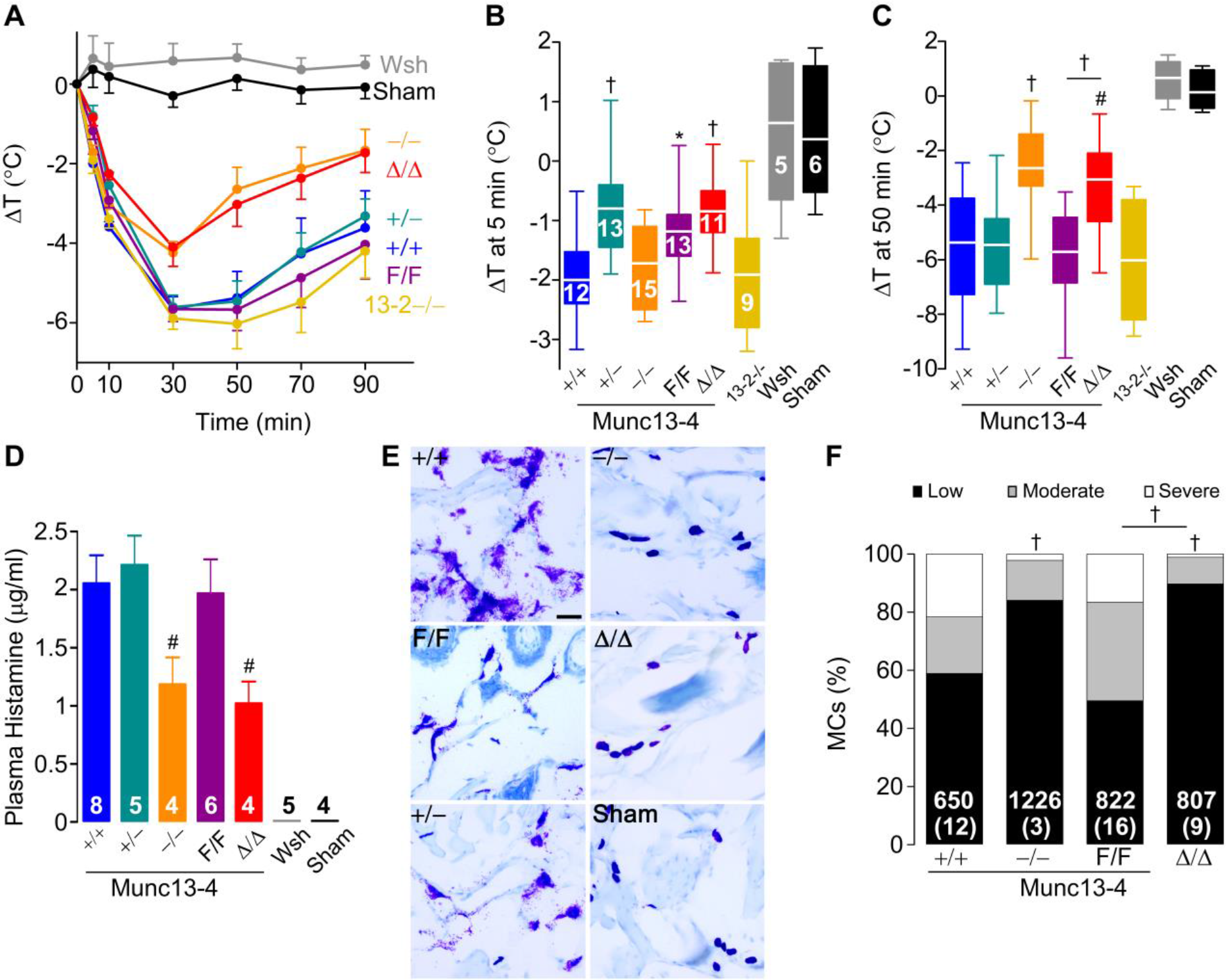
MC Munc13-4 is required for a full anaphylactic response. Mice sensitized i.v. with anti-DNP IgE and challenged i.v. with DNP-HSA. Sham, non-sensitized but challenged B6 mice. *A*, change in body core temperature from baseline (ΔT). Circle, mean; error bar, SEM. ΔT 5 minutes (*B*) and 50 minutes (*C*) post-challenge. Line, mean; box, 25^th^-75^th^ percentile; whiskers, 5^th^-95^th^ percentile; N = number inside boxes, applies to *A-C*. *D*, plasma histamine levels 3 minutes post-challenge. Bar, mean; error bar, SEM; N = number inside bars. *E*, representative lip sections 15 minutes post-challenge stained with toluidine blue to identify MC metachromatic granules. Scale bar = 40 μm. *F*, quantification of MC degranulation in lips and tongues based on proportion of granules outside the MC: low < 10%, moderate 10-50% and severe > 50%. Number of cells inside bars, number of animals in parentheses. # = p < 0.05, † = p < 0.01, * = p < 0.001; all comparisons are to Munc13-4^+/+^ unless otherwise indicated.

During anaphylaxis, MC proteases contained in MC granules can be detected in serum (36). We measured plasma histamine, also stored in MC granules (37), as a surrogate of MC degranulation in mice undergoing anaphylaxis. We took special care during plasma collection to prevent the release of histamine from platelets. We found decreased plasma levels of histamine in challenged Munc13-4^−/−^ and Munc13-4^Δ/Δ^ mice (Fig. 2*D*). In histological samples of connective tissues from challenged Munc13-4^+/+^, Munc13-4^+/−^ and Munc13-4^F/F^ mice, we observed multiple metachromatic granules surrounding degranulated MCs. Conversely, most visible MC granules from challenged Munc13-4^−/−^ and Munc13-4^Δ/Δ^ mice remained densely packed inside the MCs, which were almost indistinguishable from those of unchallenged Munc13-4^+/+^ mice (Fig. 2*E*). When quantified with a grading system (supplemental Fig. S2), we observed almost no MCs that we would grade as severely degranulated in the absence of Munc13-4 (Fig. 2*F*). There was a clear correlation between Munc13-4 expression in MCs, the severity of MC degranulation, the histamine plasma levels and the hypothermic response to anaphylaxis.

### Characterization of MCs from Munc13-4 mutant mice

Abnormalities in the number, distribution and development of MCs caused by Munc13-4 deficiency could also blunt the anaphylactic response. Consequently, we studied the baseline characteristics of the mutant mice and their MCs. As shown in Figure 3 and Table 1, we found no differences in the density of MCs in the dermis, the distribution of MCs in other tissues (not shown), and the proportion and absolute number of MCs in peritoneal lavages. Qualitatively, the metachromasia of granules from all MCs was almost identical. The same proportion and number of bone marrow cells underwent differentiation into BMMCs in media enriched with stem cell factor (SCF) and IL3. Finally, all MCs and BMMCs expressed similar levels of receptors important for MC development and activation on their surfaces (FcεRIα, Kit and IL33R). We assessed cell morphology by stereology of EM profiles of peritoneal MCs fixed immediately after harvesting. Because the area of the cell profiles (A) and cell surface density (Sv cell) were similar, we concluded that there was no difference in the size, shape and surface complexity of the MCs. The fact that both granule volume density (Vv) and granule surface density (Sv granule) did not differ indicated that the number, size and shape of the MC secretory granules at baseline is unchanged. Therefore, we found no baseline abnormality that could explain the phenotype of Munc13-4^−/−^ and Munc13-4^Δ/Δ^ mice, or that could interfere with the interpretation of the cell assays described below.

**Figure 3.**
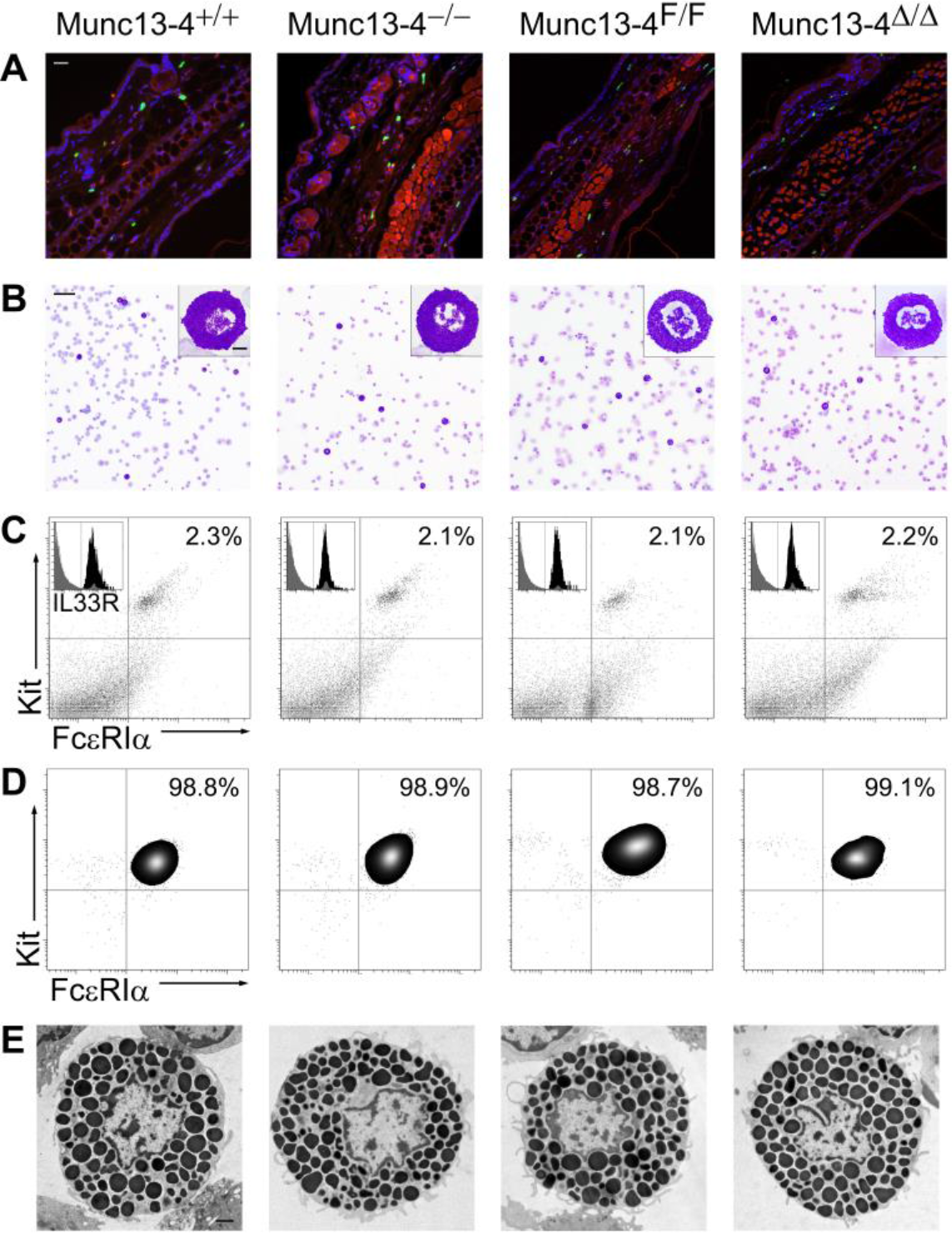
Characterization of MCs from Munc13-4 mutant mice. Representative samples; for detailed quantification please refer to Table 1. *A*, ear sections stained with FITC-avidin (green) and Hoechst (blue). MCs are identified as FITC^+^ granules surrounding Hoechst^+^ nuclei; cartilage and epithelial autofluorescence in the red channel delimited the dermis. Scale bar = 50 μm. *B*, Wright-Giemsa staining of peritoneal lavages. MCs are identified by their metachromatic granules. Scale bar = 80 μm, inset scale bar = 5 μm. *C*, flow cytometry of peritoneal lavage cells labeled for Kit and FcεRIα. MCs are identified as Kit^+^/FcεRIα^+^ double-positive cells. Inset: Kit+/FcεRIα^+^ double-positive cells before (grey) and after (black) adding labeled anti-IL33R antibody. *D*, flow cytometry of BMMCs after 6 weeks in media enriched with SCF and IL3 labeled for Kit and FcεRIα antibodies. *E*, EM cell profiles of peritoneal MCs used for stereology. Scale bar = 2 μm.

**Table 1.** Characterization of MCs from Munc13-4 mutant mice^£^

|  | **Munc13-4<sup>+/+</sup>** | **Munc13-4<sup>-/-</sup>** | **Munc13-4<sup>F/F</sup>** | **Munc13-4<sup>Δ/Δ</sup>** |
| --- | --- | --- | --- | --- |
| **Dermal MCs** |  |  |  |  |
| Density (MCs/mm<sup>2</sup> of dermis)* | 220.9 ± 20 | 204.9.7 ± 42 | 210.2 ± 17.6 | 209.3 ± 20.9 |
| **Peritoneal MCs** |  |  |  |  |
| Count (MCs/μl of lavage)† | 23.67 ± 6 | 23.9 ± 6.7 | 23.3 ± 1.4 | 23.07 ± 12 |
| MCs (% of lavaged cells)† | 3.35 ± 0.6 | 3.07 ± 0.4 | 2.98 ± 0.7 | 3.1 ± 1.3 |
| V<sub>v</sub>‡ | 0.51 ± 0.01 | 0.52 ± 0.01 | 0.54 ± 0.01 | 0.53 ± 0.03 |
| S<sub>v</sub> cell (μm<sup>-1</sup>)‡ | 0.49 ± 0.02 | 0.50 ± 0.03 | 0.49 ± 0.02 | 0.49 ± 0.04 |
| S<sub>v</sub> granule (μm<sup>-1</sup>)‡ | 6.98 ± 0.1 | 7.05 ± 0.2 | 7.01 ± 0.2 | 6.97 ± 0.4 |
| A (μm<sup>2</sup>)‡ | 84.61 ± 2 | 85.32 ± 1.8 | 83.75 ± 2.3 | 85.76 ± 2.4 |
| Kit<sup>+</sup>/FcεRIα<sup>+</sup> (% of total) | 2.2 ± 0.2 | 1.9 ± 0.3 | 2.1 ± 0.5 | 2.2 ± 0.3 |
| IL33R (% of DP)§ Φ | 98.3 ± 0.3 | 97.1 ± 0.2 | 98.5 ± 0.3 | 97.6 ± 1.0 |
| Kit MFI (AU)¶ | 7.3 ± 0.8 | 8.4 ± 0.6 | 7.4 ± 0.8 | 8.7 ± 0.5 |
| FcεRIα MFI (AU)¶ | 5.0 ± 0.5 | 5.0 ± 0.5 | 4.4 ± 0.4 | 4.9 ± 0.7 |
| IL33R MFI (AU)¶ | 8.2 ± 0.8 | 7.5 ± 0.3 | 7.6 ± 0.8 | 8.3 ± 1.1 |
| **BMMCs** |  |  |  |  |
| Kit<sup>+</sup>/FcεRIα<sup>+</sup> (%)§ | 97.9 ± 0.4 | 96.8 ± 1.0 | 95.2 ± 1.8 | 96.0 ± 1.2 |
| Kit MFI (AU)¶ | 5.0 ± 0.6 | 5.2 ± 0.6 | 5.2 ± 0.5 | 4.8 ± 0.8 |
| FcεRIα MFI (AU)¶ | 4.8 ± 0.3 | 5.1 ± 0.5 | 5.4 ± 0.5 | 4.7 ± 0.7 |

<sup>‡</sup>Results are means ± SEM for 10 mice of each genotype. A, area; AU, arbitrary units; BMMC, bone marrow-derived MC; DP, double positive cells for Kit and FcεRIα; IL33R, receptor for interleukin 33; MC, mast cell; MFI, mean fluorescence intensity; S<sub>v</sub>, surface density; V<sub>v</sub>, volume density.
\*cells with FITC<sup>+</sup> granules and a Hoeschst<sup>+</sup> nucleus per mm<sup>2</sup> of dermis in random ear sections (10 sections/mouse).
†Neubauer chamber counts and differentials of cytopspins of peritoneal lavages.
‡Stereological analysis of randomly acquired EM images of peritoneal MCs (10 cell profiles/mouse).
§Fraction of Kit<sup>+</sup>/FcεRIα<sup>+</sup> double-positive cells by flow cytometry.
ΦFraction of Kit<sup>+</sup>/FcεRIα<sup>+</sup> double-positive cells that were also IL33R positive.
¶Cell surface expression of Kit, FcεRIα and IL33R expressed as MFI of specific labelled primary antibodies.

### Lack of Munc13-4 affects exclusively MC regulated exocytosis

The defective MC degranulation in mice lacking Munc13-4 prompted us to address whether all effector responses were compromised by measuring the secretion of products that putatively depend on regulated exocytosis, constitutive exocytosis and non-exocytic transport. PCMCs were stimulated via FcεRIα or by a combination of the DAG-analog phorbol 12-myristate 13-acetate (PMA) and the Ca^2+^ ionophore ionomycin. We quantified the fraction of total cell histamine and β-hexosaminidase secreted after stimulation. These two compounds are preformed, stored in MC secretory granules, and secreted via regulated exocytosis. In both cases, we detected the expected bell-shaped response to increasing amounts of IgE-ligand (38,39) in Munc13-4^+/+^ PCMCs, while PCMCs from Munc13-4^−/−^ and Munc13-4^Δ/Δ^ mice had a flat response (Fig. 4, *A* and *B*). We could detect a small response in the hypomorphic Munc13-4^F/F^ PCMCs, which was magnified when we used PMA/ionomycin (PI) as agonist (Fig. 4*C*). The almost identical results between Munc13-4^−/−^ and Munc13-4^Δ/Δ^ PCMCs points to the efficiency of Cre recombination in these cells.

**Figure 4.**
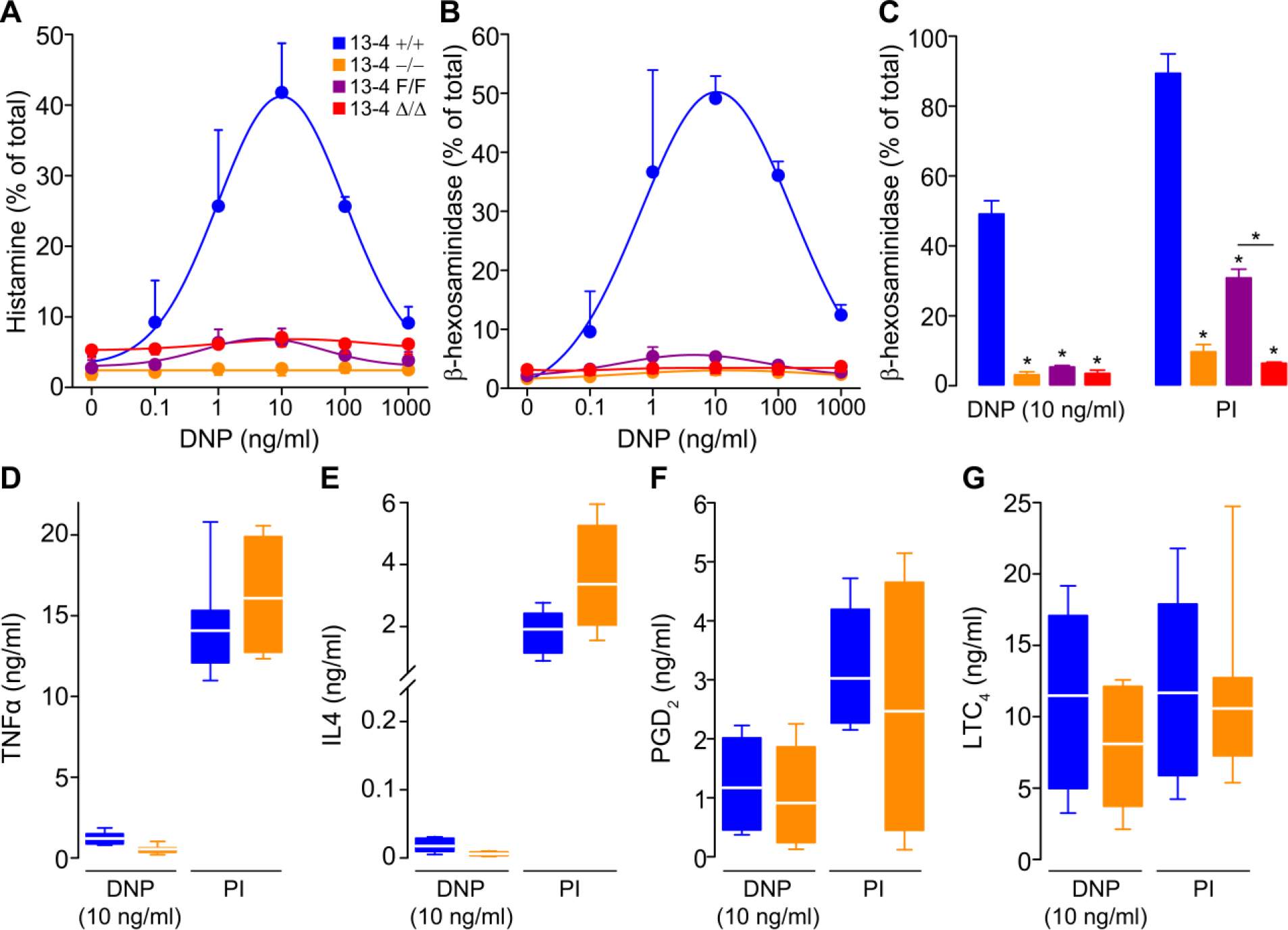
Munc13-4 controls only regulated exocytosis in MCs. PCMCs sensitized with anti-DNP IgE and then challenged with DNP-HSA (DNP) or PMA/ionomycin (PI). Color legend in *A* applies to all panels. Regulated exocytosis was monitored by measuring secretion of histamine (*A*) or β-hexosaminidase (*B* and *C*) as fraction of total cell content 30 minutes post-stimulation. N = 4; Circle and bar, mean; error bar, SEM. Constitutive exocytosis was monitored by measuring secretion of TNFα (*D*) and IL4 (*E*) 6 hours poststimulation. N = 8. Secretion independent of exocytosis was monitored by measuring PGD_2_ (*F*) and LTC_4_ (*G*) 30 minutes post-stimulation. N =4. Line, mean; box, 25^th^-75^th^ percentile; whiskers, 5^th^-95^th^ percentile.* = p < 0.001; all comparisons are to Munc13-4^+/+^ unless otherwise indicated.

We chose the dose of DNP at which we detected the largest response (10 ng/ml) and PI to interrogate other MC effector responses. To address constitutive exocytosis of newly synthesized cytokines, we measured the secretion of TNFα and IL4, and for responses independent of exocytosis, we measured the secretion of PGD_2_ and LTC_4_. As reported by others, we found that engaging IgE signaling was a weaker stimulus than other agonist for cytokine production (40). In either case, we found no differences between Munc13-4-sufficient and Munc13-4-deficient MCs (Fig. 4, *D* and E). Also, we observed no differences in secretion of PGD_2_ and LTC_4_ (Fig. 4, *F* and *G*). TNFα, IL4, PGD_2_ and LTC_4_ were undetectable in supernatants of unstimulated cells (not shown). Thus, we found that only regulated exocytosis is affected by the absence of Munc13-4.

### Munc13-2 and Munc13-4 regulate the kinetics of MC exocytosis

Based on our findings for regulated exocytosis and anaphylaxis, we decided to study the influence of Munc13 proteins on the dynamics of exocytosis at the single-cell level. To detect exocytosis in individual MCs, we measured changes in membrane capacitance in the whole-cell patch clamp recording configuration. Intracellular dialysis of GTPγS and Ca^2+^ through the patch pipette induces almost complete MC degranulation, and the resolution of this technique is such that we can record individual granule-to-plasma membrane fusion events (41–44). Membrane capacitance (C_m_) is proportional to the area of the plasma membrane, which increases by the addition of membrane from a secretory vesicle upon fusion, and is recorded as an almost instantaneous “step up” in capacitance gain (ΔC_m_) (43). We tested freshly isolated peritoneal MCs from all Munc13-4 genotypes, Munc13-2 KO mice and double Munc13-2/Munc13-4 KO mice (DKO). As a negative control we used Munc13-4^+/+^ MCs penetrated with a pipette loaded with an intracellular solution lacking GTPγS to demonstrate that the manipulation itself was not responsible for the changes in C_m_ (Fig. 5*A*) The baseline C_m_ of all the MCs studied was 6.1 ± 0.4 pF, the intracellular Ca^2+^ concentration achieved in all MCs as measured by ratiometry was 701 ± 44 nM (mean ± SEM in both cases), and we found no differences in these values among all the genoptypes. We recorded the ΔC_m_ over this baseline after stimulation to quantify the total amount of exocytosis (Fig. 5, *A* and *B*). We then normalized the C_m_ curves (Fig. 5*C*) to obtain the rate of gain between 40% and 60% of total ΔC_m_, which corresponds to the steepest part of each curve (Fig. 5*D*). We used the C_m_ traces (Fig. 6*A*) to measure the interval between the beginning of stimulation and the beginning of exocytosis in each MC (Fig. 6*B*). Finally, we measured the sizes of steps in C_m_ curves from Munc13-4-sufficient and Munc14-3-deficient MCs (Fig. 6, *C* and *D*) to estimate the size of the vesicles being exocytosed.

Munc13-4 deficiency (Munc13-4^−/−^ and Munc13-4^Δ/Δ^) resulted in severe reductions in the total amount of exocytosis. Any residual exocytosis was very slow and delayed. The mutants with intermediate expression levels of Munc13-4 (Munc13-4^+/−^ and Munc13-4^F/F^) had a normal interval to start of exocytosis, and they reached a normal level of ΔC_m_ but at a slow rate. Indeed, we found a clear dose-dependency between the expression levels of Munc13-4 and the rate of exocytosis in MCs. The frequency distribution of the step sizes was very different between Munc13-4^+/+^ and Munc13-4^−/−^ MCs, with a clear shift to the left and absence of large fusion events in the absence of Munc13-4.

**Figure 5.**
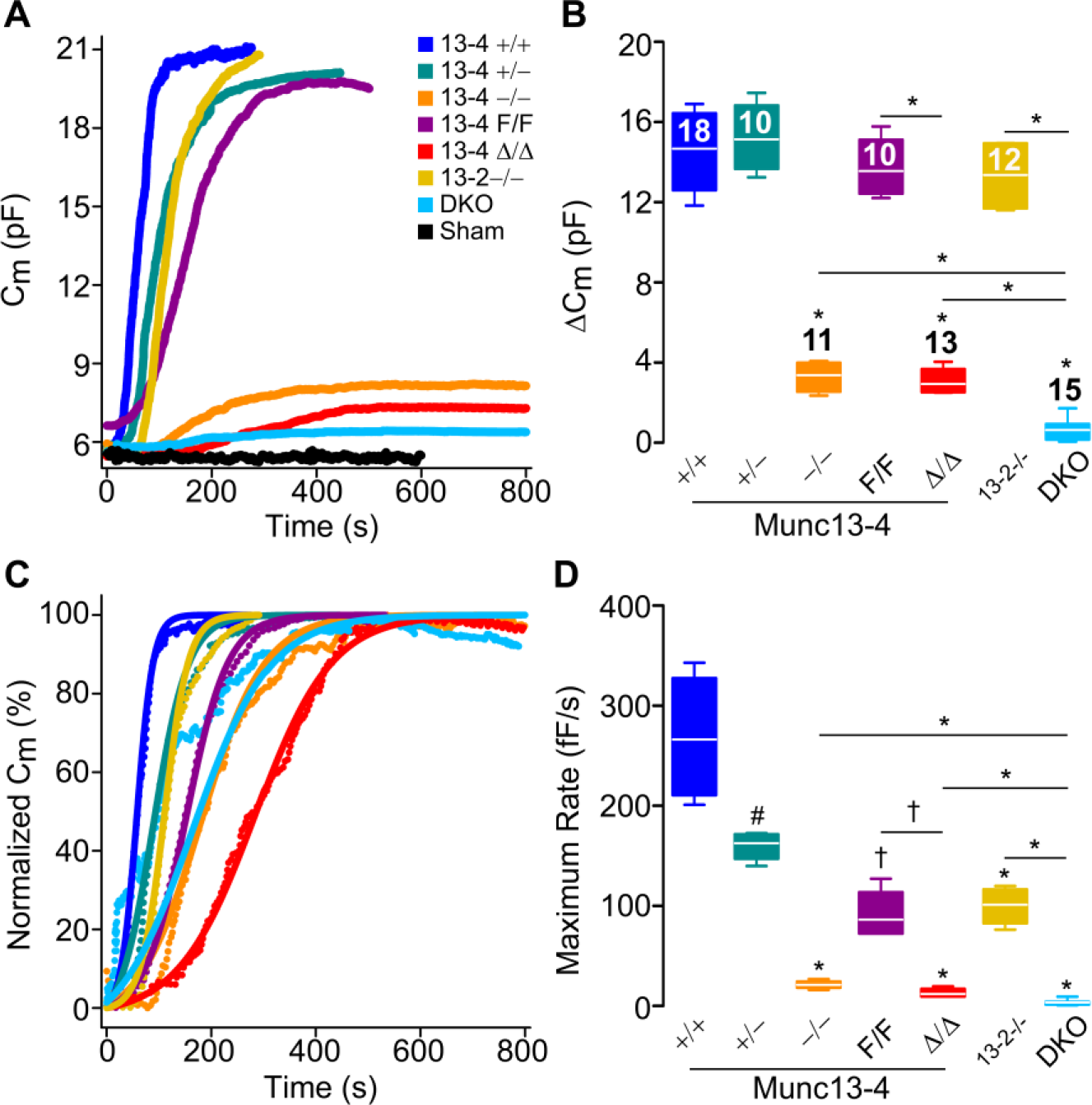
Munc13-4 regulates the amount and rate of MC exocytosis. Capacitance (C_m_) recordings from peritoneal MCs during exocytosis stimulated by intracellular dialysis of GTPγS and Ca^2+^. Access to the cell interior under the whole-cell patch clamp recording technique was achieved at Time = 0. Sham, Munc13-4^+/+^ MCs dialyzed with a solution lacking GTPγS. Color legend in *A* applies to all panels. *A*, representative traces of cumulative C_m_. *B*, total C_m_ change (ΔC_m_) above baseline. N = number inside boxes, applies to *B* and *D. C,* representative normalized C_m_ traces. *D*, maximum rate of ΔC_m_ (rate between 40%-60% of total ΔC_m_). Line, mean; box, 25^th^-75^th^ percentile; whiskers, 5^th^-95^th^ percentile. # = p < 0.05, † = p < 0.01, * = p < 0.001; all comparisons are to Munc13-4^+/+^ unless otherwise indicated.

**Figure 6.**
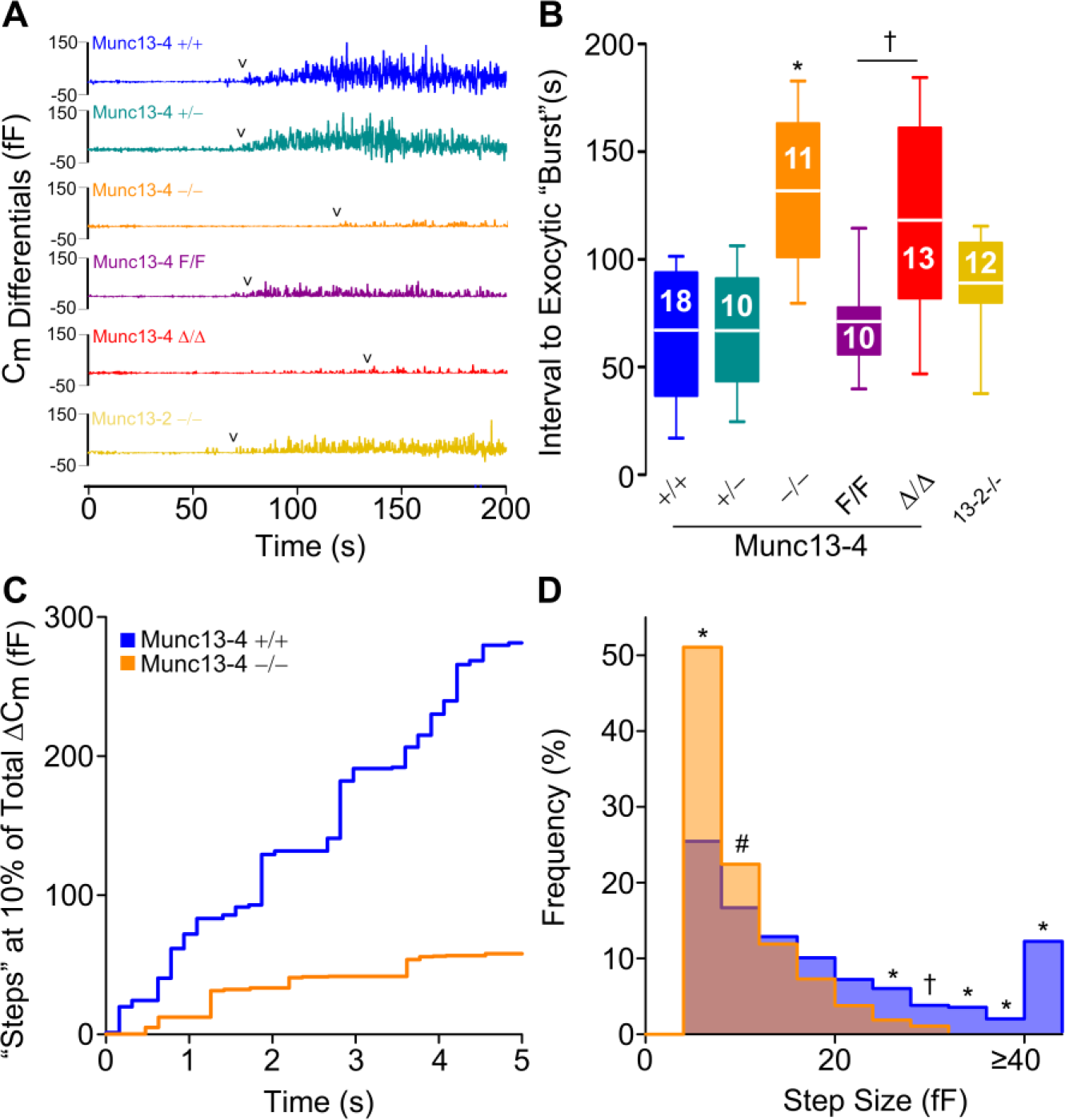
Munc13-4 regulates the start of exocytosis and the size of the “steps” in C_m_. *A*, representative C_m_ differentials from C_m_ traces shown in Fig. 5*A*. 0, time of cell access; arrowheads, first C_m_ differential ≥ 8 fF. Color legend applies to *A* and *B*. *B*, interval between start of dialysis of GTPγS and Ca^2+^ and beginning of the exocytic “burst”. Line, mean; box, 25^th^-75^th^ percentile; whiskers, 5^th^-95^th^ percentile; N = number inside boxes. *C*, representative Cm traces at ~10% of total ΔC_m_. Signals < 0 fF have been removed for clarity. Color legend applies to *C* and *D*. *D*, histogram of C_m_ step sizes between 1% and 15% of total ΔC_m_. Signals < 4 fF have been removed for clarity. N = numbers in *B*, ~68 steps/cell. # = p < 0.05, † = p < 0.01, * = p < 0.001; all comparisons are to Munc13-4^+/+^ unless otherwise indicated.

Although deletion of Munc13-2 in isolation had no effects on the total amount of exocytosis, it partially impacted the rate of exocytosis, and any residual exocytosis we observed in Munc13-4-deficient MCs was almost completely eliminated when both Munc13-2 and Munc13-4 were deleted. The defect was so severe in the DKO MCs that we could not determine a start point of exocytosis or have enough steps to analyze. Therefore, the amount, rate, interval between stimulation and start of exocytosis, and size of the vesicular compartments fusing with the plasma membrane, all depend on Munc13-4, and a role for Munc13-2 was revealed in the absence of Munc13-4.

### Lack of Munc13-4 alters the ultrastructural changes associated with exocytosis in MCs

A limitation of patch clamp is that only events at the cell surface generate a signal. Consequently, it cannot distinguish between individual vesicles fusing with the plasma membrane and sequential compound exocytosis connecting the membrane to inward granules. Also, the lack of large steps in ΔC_m_ could reflect a failure of homotypic granule-to-granule fusion, of heterotypic fusion of large multigranular compartments to the plasma membrane, or both. To overcome this deficit, we studied stimulated peritoneal MCs under EM and stereology. While there were marked changes between stimulated and unstimulated Munc13-4^+/+^ MCs, stimulated Munc13-4^−/−^ MCs were almost indistinguishable from unstimulated controls (Fig. 7*A*). To quantify these qualitative differences we based our stereological analysis on EM signs of MC activation: the granules increase in size as they become hydrated and they lose electron-density as the contents become diluted (45), the geometry of the surface of the cell increases in complexity as membrane is added during degranulation, and there is loss of the intracellular boundaries between granules as they fuse during compound exocytosis (42,46). We show that while the cell profiles of stimulated Munc13-4^+/+^ MCs had a decrease in compactness of their granules, displayed as a right shift in relative electron-lucency (Fig. 7*B*, blue area), the density of the granules in Munc13-4^−/−^ cell profiles was basically unchanged (red area) compared to unstimulated controls (black area). This was confirmed when we measured the Vv of granules with a relative electron-lucency > 150 (7*C*). As the diameter of a sphere increases, its surface-to-volume ratio (expressed by the surface density or Sv) decreases, so the reduction in granule Sv in stimulated WT cells compared to unstimulated WT cells reflects the swelling of MC granules during activation, which did not occur in stimulated Munc13-4^−/−^ MCs (Fig. 7*D*). The cell Sv increases as the surface complexity increases and the cell becomes less spherical, and again the stimulated Munc13-4^−/−^ MCs behaved more like unstimulated Munc13-4^+/+^ cells than stimulated Munc13-4^+/+^ cells (Fig. 7*E*). Finally, we quantified the fraction of multigranular complexes sharing a single membrane (supplemental Fig. S3) and observed that, despite stimulation, none could be detected in the absence of Munc13-4 (Fig. 7*F*). Thus, Munc13-4 not only controls the heterotypic fusion of secretory granules with the plasma membrane, it also mediates the homotypic fusion between granules.

**Figure 7.**
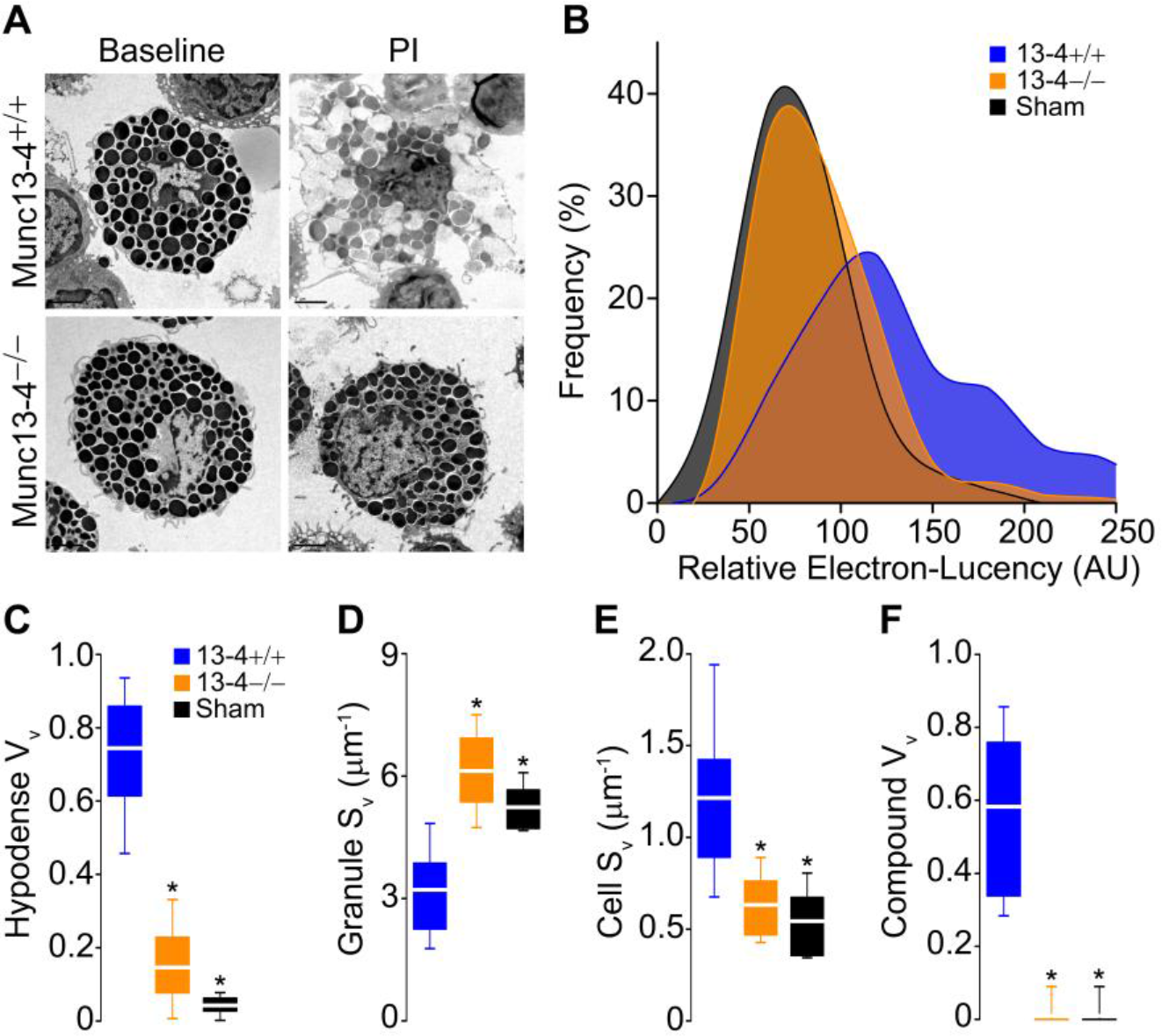
Lack of Munc13-4 alters the ultrastructural changes associated with MC exocytosis. Peritoneal MCs fixed after exposure to PMA/ionomycin (PI). Sham, Munc13-4^+/+^ MCs exposed to vehicle. *A*, representative cell profiles before and after stimulation. Scale bar = 4 μm. *B*, relative electron-lucency of MC granules based on a 0 (black) to 255 (white) scale. N = ~40 granules/cell, 30 cells/animal, 4 animals. *C-F*, stereological analysis of MC profiles. Vv, volume density; Sv, surface density; hypodense, granules with relative electron-lucency > 150; compound, multigranular profiles sharing a single delimiting membrane. N = 30 cells/animal, 4 animals. * = p < 0.001; all comparisons are to Munc13-4^+/+^ unless otherwise indicated.

## DISCUSSION

MCs store biogenic amines, lysosomal enzymes, proteases and highly sulfated glycosaminoglycans attached to a serglycin protein backbone in their large secretory granules (29,47), and the local and systemic release of these substances contributes to many pathologic processes (48–51).

Munc13-4 has been detected in BMMCs (52,53) and RBL-2H3 cells (26), and we found that Munc13-4 is co-expressed with Munc13-2 in mature MCs (Fig. 1). Munc13-2 has two splice variants, one expressed predominantly in brain (bMunc13-2) and another expressed ubiquitously (ubMunc13-2) (12,54). ubMunc13-2 is the variant expressed in MCs, and its structure is very similar to that of Munc13-1 (55). The Munc13-2 KO mouse line we used lacks both splice variants (56). With this deletant mouse, we were able to detect a role for Munc13-2 in MC exocytosis (Fig. 5). It is possible that a subpopulation of MC secretory granules uses Munc13-2 for priming, and that this could only be revealed in a Munc13-4-null background. Another explanation is that, similar to cytotoxic T lymphocytes where expression of Munc13-1 can supplement or supplant the function of Munc13-4 in exocytosis of lytic granules (57), Munc13-2 might partially replace the function of Munc13-4 in exocytosis of MC granules. Our findings in electrophysiology and EM that there was almost no residual compound multigranular exocytosis once Munc13-4 was removed (Figs. 6*D* and 7*F*), indicate that not all Munc13-4 functions can be replaced by Munc13-2.

While developing the Munc13-4^F/F^ line, we disrupted the expression of Munc13-4, creating a hypomorphic mouse with expression levels between that of heterozygous and homozygous deletants (Fig. 1). In all assays where we compared Munc13-4^F/F^ with Munc13-4^Δ/Δ^ mice we found a significant difference, indicating that this conditional KO line is a very useful loss-of-function model. Even more, we took advantage of this unexpected outcome and we found a dose-dependent relationship between MC exocytosis and expression levels of Munc13-4 (Figs. 4 and 5).

By C_m_ measurements, we detected a 78% decrease in stimulated MC exocytosis in the absence of Munc13-4 (Fig. 5*B*). This defect was not due to differences in MC granules (Fig. 3 and Table 1), as is the case in other secretion anomalies (58,59). The defect in ΔC_m_ we found in Munc13-4-deficient mice was confirmed by the almost complete lack of evidence of degranulation in our EM stereology studies (Fig. 7). The close correlation between our electrophysiological and morphological studies indicates that stereology of EM cell profiles is a reliable method to assess exocytosis in MCs. Although a participation of Munc13-4 in exocytosis from BMMCs (52) and RBL-2H3 cells (25,26,60) has been described before, our findings in MCs with intermediate expression of Munc13-4 (+/− and F/F) proves that Munc13-4 is a rate-limiting factor in this process (Figs. 4*C* and 5*D*).

In most cells, only a fraction of their secretory vesicles are primed at rest, and this pool increases as more Munc13 is recruited to the plasma membrane during cell activation (61). A defect in priming could explain why Munc13-4 is rate-limiting for MC secretion. First, the longer interval between MC stimulation and response corresponds to a delayed “burst” of degranulation that we observed in the absence of Munc13-4 (Fig. 6*B*), indicating that this protein is a limiting factor for the number of secretory granules that may be fusion-competent before MC activation, similar to the ready releasable pool (RRP) of synaptic vesicles in neurons (62). Second, Munc13-4 is required for the acceleration of exocytosis after activation (Fig. 5*D*), indicating that Munc13-4 might be a limiting factor for the recruitment of new secretory vesicles to the primed pool, similar to the replenishment of the RRP in neurons (63). Despite these similarities to Munc13-1-dependent neuronal priming, Munc13-4 lacks the regulatory C2A and C1 domains, implying that homodimerization, interactions with RIM and binding to DAG are dispensable for these Munc13-4 functions (17,18,20,24).

The close correlation between the maximum rate of exocytosis and the level of expression of Munc13-4 we observed (Fig. 5*D*) could have another explanation. Mechanistically, the transition from initial to maximum rate of exocytosis in MCs could be achieved by compound exocytosis (42). The domino-like fusion of individual granules to other granules that have already fused with the plasma membrane in sequential compound exocytosis decreases the distance between vesicle and target membranes, accelerating the rate of single-vesicle fusion. Also, the fusion between granules to create large secretory compartments before fusing with the plasma membrane in multigranular compound exocytosis adds a larger membrane surface per exocytic event. The membrane of a single MC granule has a C_m_ of approximately 7 fF (64). We found that most fusion events in stimulated Munc13-4^+/+^ MCs are larger than 8 fF, indicating that multigranular compound exocytosis is common in normal MCs. In contrast, most steps recorded in Munc13-4^−/−^ MCs were ≤ 8 fF (Fig. 6*D*), implying that most exocytosis remaining in the absence of Munc13-4 is of single vesicles. The lack of large “steps” could be due to a failure to form large multigranular compartments or a defect in fusion of these multigranular compartments with the plasma membrane. Our EM stereology studies of activated peritoneal MCs confirmed the former (Fig 7*F*). This loss-of-function model and the combination of electrophysiology and EM stereology corroborate the long suspected dependency of the accelerated phase of MC exocytosis on compound multigranular exocytosis.

Our findings prove that Munc13-4 is required for both the heterotypic fusion of granule membranes with the plasma membrane and the homotypic fusion of granule membranes in mature MCs. Others have reported that the two C2 domains of Munc13-4 are required for the homotypic fusion responsible for the formation of large vacuoles in RBL-2H3 cells (26). The fact that both types of fusion are under the control of the same Munc13 protein suggests that both membrane fusion machineries may share other fundamental components.

Using populations of MCs and other stimuli we found no difference in secretion of PGD_2_ and LTC4 between Munc13-4^+/+^ and Munc13-4^−/−^ MCs. This shows that Munc13-4 does not play a role in these exocytic-independent MC effector responses, that Munc13-4-deficient MCs can be activated, and that we are not dealing with a failure in signaling but with a pure exocytic defect. Munc13-4-deficient MCs had a dose-dependent defect in the secretion of preformed mediators stored in MC secretory granules (Fig. 4) that could not be explained by changes in size, number or shape of MC granules (Fig. 3 and Table 1). In many systems, the presence of a synaptotagmin and a Munc13 protein defines an exocytic process as regulated (10). Similar to what we found in the synaptotagmin-2 KO mouse (38), deficiency of Munc13-4 affected only the release of MC granules, lending further support to the concept that this process is a form of regulated exocytosis.

Our manipulations of the expression of Munc13-4 did not affect the secretion of cytokines by MCs, as was the case for synaptotagmin-2 and VAMP-8 (38,65), pointing to the existence of an alternative pathway for cytokine secretion. This is supported by studies in which MCs could be stimulated to secrete cytokines without inducing degranulation (66,67). The most parsimonious explanation is that de novo generated mediators are secreted via constitutive exocytosis using a distinct secretory route (68–70), and that this process is not controlled by Munc13-4. Our finding that the densities of receptors on the MC surface were the same regardless of the expression levels of Munc13-4 (Fig. 3 and Table 1) supports this possibility, given that translocation of most receptors to the plasma membrane depends on constitutive exocytosis (2). Others have found TNFα inside MC granules (71), and it has been documented that vesicles containing TNFα can merge with MC secretory granules (72,73) in a process that also requires Munc13-4 (26), suggesting that this cytokine uses regulated exocytosis as the final step in its secretion. Unfortunately, none of these studies addressed the relative contribution of this putative secretory pathway to the overall secretion of TNFα by MCs. A way to reconcile these results with our findings of normal TNFα secretion (Fig. 4*D*) despite a severe defect in degranulation in the absence of Munc13-4 is that the contribution of degranulation to TNFα secretion is minuscule. The existence of a discrete subpopulation of MC granules that contain TNFα but that is not controlled by Munc 13-4 is incompatible with our results, unless these granules also lack β-hexosaminidase and histamine.

We did not find that the decreased or absent expression of Munc13-4 caused any alteration in the number, distribution and differentiation of MCs in the mutant mice (Fig. 3 and Table 1). Thus, any abnormality in the animals should reflect exclusively a functional defect in MCs. There was no overall manifestation of haploinsufficiency in our anaphylaxis model. Even the hypomorphic Munc13-4^F/F^ mice reached the same degree of hypothermia at 50 minutes, increases in circulating histamine and histological evidence of MC degranulation in connective tissues compared to Munc13-4^+/+^ mice (Fig. 2), indicating that these manifestations depend not on the rate but on the total amount of exocytosis in MCs (Fig. 5). The only significant difference was detected in the early time points (5 minutes after challenge), when Munc13-4^+/−^ and Munc13-4^F/F^ were significantly less hypothermic than Munc13-4^+/+^ mice. This correlates with the slower rates of exocytosis we found in the electrophysiological studies of MCs from these two mutants. These observations indicate that the earliest response engaged is MC degranulation, making the immediate anaphylactic response very sensitive to changes in the kinetics of MC regulated exocytosis. Afterwards, as the reaction progresses and MCs achieve full degranulation, and engage other effector responses, a partial deficiency in Munc13-4 has no effects.

We found a correlation between less MC degranulation, less plasma histamine and less hypothermia (Fig. 2). Because the responses in Munc13-4^−/−^ and Munc13-4^Δ/Δ^ mice were indistinguishable, we conclude that the Munc13-4-dependency of this response is limited to MCs, and that the failure in MC degranulation *in vivo* was not contingent to a Munc13-4-dependent abnormality in another cell that indirectly affected MC activation. The protection was significant but not absolute. Cma1/mMCP5 is expressed exclusively in MCs, but not in all MCs; it is expressed mainly in constitutive/connective tissue MCs (74). Hence, a population of inducible/mucosal MCs that do not express Cre, and therefore do not undergo recombination in Munc13-4^Δ/Δ^ mice, could be responsible for the residual anaphylactic response. However, that does not explain why Munc13-4^−/−^ mice, which lack Munc13-4 in all cells, including mucosal MCs, have the same degree of hypothermia and levels of residual circulating histamine as Munc13-4^Δ/Δ^ mice. The possibility that cells other than MCs (e.g., basophils and platelets) are responsible for the residual response is improbable, given that MC-deficient animals, which have intact numbers of other cells important in anaphylaxis (75), exhibit complete protection from hypothermia and had undetectable levels of circulating histamine. Another explanation is that the residual exocytosis we documented in our electrophysiological and cell secretion assays (Figs. 4*A-C* and 5B) allows enough mediators to be secreted to precipitate a limited response in Munc13-4^−/−^ and Munc13-4^Δ/A^ mice (Fig. 2). Lastly, we think that MC responses independent of regulated exocytosis are involved. The duration of our model is too short to be influenced by synthesis and secretion of cytokines but coincides with the time-course of secretion of eicosanoids by MCs, which are well-known mediators of anaphylaxis (76) and do not require Munc13-4 for secretion (Fig. 4).

MC degranulation is always cited as one of the main culprits in the generation of the anaphylactic reaction. For the first time we can quantify the relative contribution of MC degranulation in this pathologic response, and compare it to other effector responses from MCs and other cells. Based on our results, interfering with MC degranulation or blocking the effects of the multiple mediators released in this process could achieve a significant but not complete control of anaphylaxis.

## EXPERIMENTAL PROCEDURES

### Mice

All genomic portions for the Munc13-4 targeting construct were obtained from B6 mice. The 5′ homology arm (3352 bp; GRCm38:Chr11:116077882-116081234, – strand) included a portion from the 5′ flanking region to intron 2 of mouse *Unc13d*, and the 3′ homology arm (3654 bp; GRCm38:Chr11:116073258-116076912, – strand) from exon 4 to intron 15. The vector contained an FRT-flanked phosphoglucokinase promoter-driven neomycin resistance gene (PGK-Neo) upstream from the insertion fragment (995 bp;:Chr11:116076909-116077904, – strand) containing a loxP-flanked exon 3. PGK-herpes simplex virus thymidine kinase (PGK-TK) was placed downstream from the homology region (Fig. 1*B*). We electroporated the vector into 129S6:B6 ES cells, and used neomycin and gancyclovir for selection. Homologous recombinant ES cells were injected into B6(Cg)-*Tyr*^*c-2J*^/J blastocysts and implanted into pseudopregnant B6 mice. Chimeric males were crossed with B6(Cg)-*Tyr*^*c-2J*^/J females and the resulting heterozygous mutants were crossed with B6.129S4-Gt(ROSA)26Sor^tm1(FLP1)Dym^/RainJ (The Jackson Laboratory; # 009086), to remove PGK-Neo and establish our conditional KO line. This “floxed” mouse was crossed with B6.C-Tg(CMV-cre)1Cgn/J mice (The Jackson Laboratory; # 006054) that express Cre recombinase ubiquitously to delete the allele in the germ line (34) and generate our global KO line, or with Tg(Cma1-cre)ARoer mice (Dr. Axel Roers, University of Cologne) for MC-specific deletion (33). We crossed our lines with B6 mice for 10 generations. Genotypes were determined by PCR of genomic DNA with primers P1 5′GGAAAG-GTGTGTCGCCATGGTG3′, P2 5′ATCCCA-GATCAAAATGCTCCCAC3′ and P3 5′CCA-ACATAAGGCTCTCTGAAGG3′. P1 and P2 were used to differentiate between the conditional (F; 725 bp) and WT (+; 636 bp) alleles, and P2 and P3 between the globally or MC-specifically deleted (– or Δ, respectively; 415 bp) and WT (+; 1268 bp) alleles. Genomic Cre was detected using primers 5′ACAGTGGT ATT CCCGGGGAGTGT3′ and 5′GTCAGTGCGTTCAAAGGCCA3′ that give a 421 bp internal control band from the native *Cma1* gene and a 555 bp band from the transgene. Littermates +/+, +/− and −/−, and F/F and Δ/Δ were used in all experiments. We also obtained Munc13-2 KO mice (Dr. Christian Rosenmund, Charité Universitaetsmedizin) (32) and MC-deficient Kit^W-sh^/Kit^W-sh^ mice (Dr. Stephen J. Galli, Stanford University) (75). All mice were kept in a pathogen-free facility and handled in accordance with the Institutional Animal Care and Use Committee of The University of Texas MD Anderson Cancer Center.

### MC harvesting and cultures

Eight ml of PBS and 2 ml of air were injected through a 27 G needle into the mouse peritoneal cavity, aspirating the paramedian inguinal fat afterwards to seal the puncture point. After massaging the abdomen for 5 minutes the fluid was aspirated from the flanks with an 18G needle. Peritoneal cells were centrifuged (380 × g, 4 °C, 5 minutes) and processed for different studies. For PCMCs cells were resuspended in 10 ml of media (IMDM with glutamine, 10% FBS, 500 U/ml penicillin, 500 U/ml streptomycin, 1X vitamins, 1X non-essential aminoacids, 100 μM Na-pyruvate, 50 nM 2-mercaptoethanol; GIBCO) with IL3 (5 ng/ml) and SCF (50 ng/ml; both from R&D), plated on 100 mm cell culture plates, and incubated for 2 weeks (37 °C, 5% CO_2_, bi-weekly media changes). For BMMCs, we flushed out the bone marrow from the diaphyses of freshly dissected tibiae and femora with 10 ml of medium (RPMI 1640, 10% FBS, 500 U/ml penicillin, 500 U/ml streptomycin, 100 μM HEPES pH 7.3, 1 mM Na-pyruvate, 1X nonessential amino acids, 0.1% 2-mercaptoethanol; GIBCO) onto a 100 mm cell culture dish. IL3 and SCF were added as above and cells were incubated for 6 weeks (37 °C, 5% CO_2_, bi-weekly media changes).

### Histology

Peritoneal MC absolute and relative numbers were determined with a Neubauer chamber and differentials of cytospins stained with modified Wright-Giemsa. Ears were excised, fixed overnight at 4 °C in 4% paraformaldehyde and embedded in paraffin. Next, 5 μm sections were stained with FITC-avidin (2 μg/ml) and Hoechst (5 μg/ml). The density of FITC^+^/Hoescht^+^ cells were reported per area of dermis, which was measured as the area between the epidermis and the subcutaneous fat and/or cartilage (77).

### Flow cytometry and cell sorting

10^6^ BMMCs, PCMCs, or peritoneal lavage cells were suspended in 250 μl PBS, blocked with 0.5 μg/ml of anti-mouse CD16/CD32 (BD Biosciences), and incubated with 0.2 μg/ml of PE-anti-mouse CD117/Kit (BD Biosciences), APC-anti-mouse FcεRIα (eBioscience) or APC-anti-mouse IL33R (BD Biosciences) for 30 minutes at 4 °C (78). The cells were washed twice with PBS and resuspended in 500 μl of cold PBS for analysis in a BD LSRII flow cytometer (BD Biosciences). For FACS, we collected the FcεRIα^+^/Kit^+^ double positive cells using a BD FACSAria Fusion Cell Sorter.

### Expression analyses

For qPCR, FACS-sorted peritoneal MCs were immersed in DNA/RNA Shield (Zymoresearch) for 1 day and homogenized by vortexing for 1 minute. RNA was extracted using RNeasy Mini Kit (Qiagen), and cDNA was obtained through reverse transcription with qScript cDNA SuperMix (Quanta Biosciences). Amplification was conducted on a ViiA 7 Real-Time PCR System using PerfeCTa qPCR ToughMix (Quanta Biosciences). Expression of Munc13-1 exons 3-4 (Mm01340418_m1, LifeTechnologies), Munc13-2 exons 17-18 (Mm01351419_m1), Munc13-3 exons 3-4 (Mm00463432_m1) and Munc13-4 exons 7-8 (Mm01252625_m1) were quantified relative to that of β-actin exon 6 (Mm00607939_s1). For RACE, RNA was reverse transcribed (SuperScript II, Life Technologies) and amplified with the adaptor primer and 5′AACCACCATGGCGAC-ACACC3′, which is common to all translated Munc13-4 splice variants (GRCm38). The gel-purified product was then sequenced. For immunoblots, mouse tissues were homogenized and sonicated on ice in 2 ml of lysis buffer (150 mM NaCl, 1% NP-40, 0.5% Na-deoxycholate, 0.1% SDS, 50 mM Tris pH 8.0) with 1% protease inhibitor cocktail (Sigma-Aldrich), or 10^6^ PCMCs were sonicated in 100 μl of PBS with 1% protease inhibitor cocktail. Protein content was estimated with BCA (Pierce). Lysates were run under denaturing conditions in a 7.5% bis-acrilamide gel, transferred onto a nitrocellulose membrane (both from Bio-Rad) and incubated with a rabbit polyclonal antibody (1:4000) raised against the mouse Munc13-4 peptide N-LLESRKGDREAQ-AFVKLRRQRAKQASQHAP-C. β-actin was used as loading control.

### Secretion assays

3 × 10^4^ PCMCs were incubated overnight with 100 ng/ml SPE-7 anti-DNP IgE (Sigma-Aldrich). They were next centrifuged (300 × g, 4 °C, 10 minutes), washed three times in secretion buffer (in mM: 10 HEPES pH 7.4, 137 NaCl, 2.7 KCl, 0.4 Na_2_HPO_4_, 5.6 glucose, 1.8 CaCl_2_, 1.3 MgSO_4_), resuspended in 90 μl of secretion buffer with 0.04 % BSA (37 °C). Cells were stimulated by adding 10 μl of secretion buffer alone, with 1 ng – 10 μg of DNP-HSA, or PMA (50 ng/ml) with ionomycin (1.2 μM) (all from Sigma-Aldrich). After 30 minutes (β-hexosaminidase, histamine, LTC_4_ and PGD_2_) or 6 hours (TNFα and IL4), the stimulation was halted by placing the cells on ice. Samples were centrifuged (300 × g, 4°C, 10 minutes,) and supernatants were collected. ELISA was used to quantify histamine, PDG_2_ and LTC_4_ (Cayman Chemical), and TNFα and IL4 (R&D Biosystems). For histamine and β-hexosaminidase, a duplicate of every sample was lysed by adding 150μL of 0.1% Triton X-100 (10 minutes, 37°C) to measure total cell content. To determine β-hexosaminidase activity, the lysates or supernatants (50 μl) were incubated with 100 μl of 3.5 mg/ml p-Nitro-N-acetyl-β-D-glucosaminide (pNAG; Sigma-Aldrich; 90 minutes, 37 °C), and the reaction was stopped by adding 100 μl of 400 mM glycine (pH 10.7). Absorbance at *λ*=405 nm was recorded (reference *λ*=620 nm) and corrected for dilutions.

### Electrophysiology

Peritoneal MCs were washed twice and plated onto glass-bottomed recording chambers filled with external recording solution (in mM: 136.89 NaCl, 2.6 KCl, 2 CaCl_2_, 0.493 MgCl_2_, 0.407 MgSO_4_, 0.441 KH_2_PO_4_, 0.338 Na_2_HPO_4_, 10 HEPES and 10 glucose; pH 7.3, 310 mOsm). Whole cell recordings were performed at 21-24 °C on individual peritoneal MCs using 5-6 MΩ patch pipettes coated with a silicone elastomer (Sylgard) and filled with an internal solution that both defined intracellular Ca^2+^ and induced degranulation (79,80). The internal solution contained (in mM: 135 K-gluconate, 7 MgCl_2_, 0.2 Na_2_ATP, 0.05 Li_4_GTPγS, 2.5 EGTA, 7.5 Ca-EGTA, 0.1 Fura-2, and 10 HEPES (pH 7.21, 302 mOsm). Alternatively, a similar calculated free Ca^2+^ concentration was achieved by replacing the EGTA and Ca-EGTA concentrations listed above with 5 mM EGTA, 7.5 mM Ca-EGTA and 2 mM CaCl_2_ (MaxChelator; http://web.stanford.edu/~cpatton/maxc.html). The free Ca^2+^ concentration was determined from the ratio of emitted fluorescence at 360 and 388 nm with the use of a computer-controlled monochrometer-based photometry system (79–81) and from Ca^2+^ calibration constants determined in vitro with the use of Fura-2 Calcium Imaging Calibration Kit (0-10 mM Ca-EGTA, 50 μM Fura-2; Thermo Fisher Scientific/Molecular Probes). Measurements of C_m_ were made with an EPC-9 patch-clamp amplifier controlled by Pulse software (HEKA Electronik). An 800 Hz sinusoidal, 30 mV peak-to-peak stimulus was applied around a holding potential of -70 mV, and the resultant signal analyzed using the Lindau-Neher technique (80,81) to yield C_m_ and the membrane (G_m_) and series (G_s_) conductances. For each 100 ms sweep, the average value was recorded, yielding a temporal resolution for C_m_, G_m_, and G_s_ of ~7 Hz. Cells selected for analysis met the criteria of G_m_ ≤ 1,000 pS, G_s_ ≥ 40 nS and steady-state intracellular [Ca^2+^] ~700 ± 100 nM. From recordings of C_m_ over time we obtained the total ΔC_m_, and from normalized curves the rates of ΔC_m_ from 40%-60% of total ΔC_m_. We logged individual changes in C_m_ (C_m_ differentials). We identified the beginning of the “burst” of exocytosis as successive C_m_ differentials ≥ 8 fF sustained for > 4 seconds; this avoided baseline isolated events. Then, we recorded the time interval between cell access and the first C_m_ differential ≥ 8 fF within this 4 second region. We measured the size of the “steps” between 1% and 15% of total ΔC_m_ after eliminating any signal < 4 fF to decrease noise.

### Electron microscopy and stereology

Peritoneal MCs were resuspended at rest or after 5 minutes of activation with PI, and fixed in 0.1 M Na-cacodylate buffer (pH 7.2) containing 2% glutaraldehyde (2 hours, room temperature). They were washed twice with PBS, resuspended in 0.1 M Na-cacodylate buffer containing 1% OsO_4_, washed in ddH_2_O, pelleted and embedded in 3% low-melting temperature agarose. The agarose pellets were dehydrated through an acetone series and then embedded in EMbed-812 epoxy resin (14120 EMS). Sections of 100 nm thickness were stained with uranyl acetate and lead citrate and viewed under a Tecnai 12 transmission electron microscope. Nucleated MC profiles were randomly photographed and at least 30 profiles from each sample were assessed by stereology. Vv and Sv were obtained with randomly placed dot and line grids on the cell profiles (82,83). The cell profile area (A) was estimated selecting circumference intersections in a cycloid grid with a known perimeter value. Multigranular events were defined as granule profiles not separated by a membrane (supplemental Fig. S3). To measure the relative electron-lucency of MC granules on a grey scale, a circle-cycloid stereological grid with circles of 0.0366 μm^2^ (diameter = 30 pixels) was randomly superimposed on cell profiles, 50 random circles that fell on granules, nuclear heterochromatin and electron-lucent extracellular space were selected and their gray scale value (0-255) was recorded. The values from the heterochromatin and extracellular space were used to set a linear scale to compare the values of each randomly selected granule, providing an internal scale normalized for each photograph.

### Passive systemic anaphylaxis

Adult mice (15-27 weeks old) were sensitized i.v. with anti-DNP IgE (20 μg in 200 μL of PBS). A day later, they were challenged i.v. with DNP-HSA (1 mg in 200 μL of PBS). Core temperatures were recorded at baseline and at 5, 10, 30, 50, 70 and 90 minutes with a rectal probe. For histamine quantification we took special care not to activate platelets because they also release significant amounts of histamine. After challenge, animals were placed in an isoflurane chamber and 3 minutes later blood was drawn slowly from the inferior vena cava through a 21G needle into a 1 ml syringe loaded with 110 μl of 4% Na-citrate. An equal volume of Tyrode's buffer (0.81% NaCl, 0.02% KCl, 0.09% glucose, 0. 005% NaH_2_PO_4_, 0.1% NHCO_3_) was added to each sample, and the plasma was separated by centrifugation (380 × g, 15 minutes, room temperature) and frozen at -80°C for subsequent histamine ELISA (Cayman Biomedical). Tongues and lips of mice euthanized 15 minutes post challenge were fixed in 4% paraformaldehyde (16 h, 4 °C), placed in 30% sucrose and 30% OCT (Tissue-Tek; VWR) in PBS overnight, and then embedded in OCT. Sections of 5 μm were stained with 0.1% toluidine blue (pH 0.5). MC degranulation was blindly assessed based on the fraction of granules outside the cell and described as low (0-10%), moderate (10-50%), or severe (> 50%) (supplemental Fig. S2).

### Statistical analysis

For continuous variables, we compared the means of all groups by one-way ANOVA; if a significant difference was found we applied Tukey’s test for multiple pairwise comparisons or Dunnett’s test for multiple comparisons against the control group. For categorical data, we used Pearson’s chi-squared test or Fisher’s exact test. Significance was set at p < 0.05.

## Acknowledgements

We thank Evelyn S. Brown and Margaret Gondo (U. of Houston) for her professional assistance with the electron microscopy, and Dr. Thomas C. Südhof for the backbone of the targeting vector.

## Conflict of interest

The authors declare that they have no conflicts of interest with the contents of this article.

## Author contributions

RA conceived the study. DCSM, YP, MJT, BFD and RA generated the mutant Munc13-4 lines. MAR and EIC performed the expression studies. EMR, MAR, DCM, YP, SM, JM and RA characterized the MCs. DCM, YP, SM, KB, and EMR performed in vivo assays. MAR and YP performed the secretion assays. AJD, DSM, AIR, RH and RA performed and analyzed the electrophysiology experiments. DCM, DSM, SM, AIR, LR, ES, AT, JM, EAG, ARB and RA performed the electron microscopy, histology and stereology. RA, EMR and MAR analyzed the data and prepared the manuscript. All authors contributed to the discussion of the study.

## FOOTNOTES

This project was supported by the National Institutes of Health AI093533A, HL129795, CA016672, EY007551, EY018239, and the Cancer Prevention Research Institute of Texas RP110166.

The abbreviations used are: A, area; AU, arbitrary units; B6, C57BL6/J mouse line; BMMC, bone marrow-derived MC; C_m_, capacitance; Cma1, chymase 1; DAG, diacylglycerol; ΔC_m_, capacitance gain; ΔT, change in body core temperature; DKO, Munc13-2/Munc13-4 double KO; DNP, 2,4-dinitrophenol; DP, double positive cells for Kit and FcεRIα; HSA, human serum albumin; LTC_4_, leukotriene C_4_; KO, knockout mouse; MC, mast cell; MFI, mean fluorescence intensity; Munc13, mammalian homologue of *C. elegans* uncoordinated gene 13; Neo, neomycin phosphotranferase; ND, non-detected; PCMCs, peritoneal cell-derived MCs; PCR, polymerase chain reaction; PGD_2_, prostaglandin D_2_; PGK, phosphoglucokinase promoter; PI, PMA/ionomycin; PMA, phorbol 12-myristate 13-acetate; qPCR, quantitative polymerase chain reaction; RACE, rapid amplification of cDNA ends; RIM, Rab3 interacting molecule; RRP, ready releasable pool; SCF, stem cell factor; SEM, standard error of the mean; SNAP25, synaptosomal-associated protein 25; SNARE, soluble N-ethylmaleimide-sensitive factor activated protein receptor; Stx, Syntaxin; Sv, surface density; TK, herpes simplex virus thymidine kinase; TNFα, tumor necrosis factor α; VAMP, vesicle associated membrane protein; Vv, volume density; W^sh^, MC-deficient Kit^W-sh/W-sh^ mice.

## REFERENCES

1. Jahn, R. (2004) Principles of exocytosis and membrane fusion. Ann Ny Acad Sci 1014, 170–178

2. Burgess, T. L., and Kelly, R. B. (1987) Constitutive and regulated secretion of proteins. Annu Rev Cell Biol 3, 243–293

3. Berridge, M. J. (1984) Inositol trisphosphate and diacylglycerol as second messengers. Biochem J 220, 345–360

4. Pang, Z. P., and Sudhof, T. C. (2010) Cell biology of Ca2+-triggered exocytosis. Curr Opin Cell Biol 22, 496–505

5. Pickett, J. A., and Edwardson, J. M. (2006) Compound exocytosis: mechanisms and functional significance. Traffic 7, 109–116

6. Lorentz, A., Baumann, A., Vitte, J., and Blank, U. (2012) The SNARE Machinery in Mast Cell Secretion. Front Immunol 3, 143

7. Sollner, T., Whiteheart, S. W., Brunner, M., Erdjument-Bromage, H., Geromanos, S., Tempst, P., and Rothman, J. E. (1993) SNAP receptors implicated in vesicle targeting and fusion. Nature 362, 318–324

8. Rizo, J., and Xu, J. (2015) The Synaptic Vesicle Release Machinery. Annu Rev Biophys 44, 339–367

9. Klenchin, V. A., and Martin, T. F. (2000) Priming in exocytosis: attaining fusion-competence after vesicle docking. Biochimie 82, 399–407

10. Sudhof, T. C. (2013) Neurotransmitter release: the last millisecond in the life of a synaptic vesicle. Neuron 80, 675–690

11. Hammarlund, M., Palfreyman, M. T., Watanabe, S., Olsen, S., and Jorgensen, E. M. (2007) Open syntaxin docks synaptic vesicles. PLoS Biol 5, e198

12. Brose, N., Hofmann, K., Hata, Y., and Sudhof, T. C. (1995) Mammalian homologues of Caenorhabditis elegans unc-13 gene define novel family of C2-domain proteins. J Biol Chem 270, 25273–25280

13. Betz, A., Okamoto, M., Benseler, F., and Brose, N. (1997) Direct interaction of the rat unc-13 homologue Munc13-1 with the N terminus of syntaxin. J Biol Chem 272, 2520–2526

14. Ma, C., Li, W., Xu, Y., and Rizo, J. (2011) Munc13 mediates the transition from the closed syntaxin-Munc18 complex to the SNARE complex. Nat Struct Mol Biol 18, 542–549

15. Yang, X., Wang, S., Sheng, Y., Zhang, M., Zou, W., Wu, L., Kang, L., Rizo, J., Zhang, R., Xu, T., and Ma, C. (2015) Syntaxin opening by the MUN domain underlies the function of Munc13 in synaptic-vesicle priming. Nat Struct Mol Biol 22, 547–554

16. Lai, Y., Choi, U. B., Leitz, J., Rhee, H. J., Lee, C., Altas, B., Zhao, M., Pfuetzner, R. A., Wang, A. L., Brose, N., Rhee, J., and Brunger, A. T. (2017) Molecular Mechanisms of Synaptic Vesicle Priming by Munc13 and Munc18. Neuron 95, 591–607 e510

17. Camacho, M., Basu, J., Trimbuch, T., Chang, S., Pulido-Lozano, C., Chang, S. S., Duluvova, I., Abo-Rady, M., Rizo, J., and Rosenmund, C. (2017) Heterodimerization of Munc13 C2A domain with RIM regulates synaptic vesicle docking and priming. Nat Commun 8, 15293

18. Liu, X., Seven, A. B., Camacho, M., Esser, V., Xu, J., Trimbuch, T., Quade, B., Su, L., Ma, C., Rosenmund, C., and Rizo, J. (2016) Functional synergy between the Munc13 C-terminal C1 and C2 domains. Elife 5

19. Shin, O. H., Lu, J., Rhee, J. S., Tomchick, D. R., Pang, Z. P., Wojcik, S. M., Camacho-Perez, M., Brose, N., Machius, M., Rizo, J., Rosenmund, C., and Sudhof, T. C. (2010) Munc13 C2B domain is an activity-dependent Ca2+ regulator of synaptic exocytosis. Nat Struct Mol Biol 17, 280–288

20. Xu, J., Camacho, M., Xu, Y., Esser, V., Liu, X., Trimbuch, T., Pan, Y. Z., Ma, C., Tomchick, D. R., Rosenmund, C., and Rizo, J. (2017) Mechanistic insights into neurotransmitter release and presynaptic plasticity from the crystal structure of Munc13-1 C1C2BMUN. Elife 6

21. Michelassi, F., Liu, H., Hu, Z., and Dittman, J. S. (2017) A C1-C2 Module in Munc13 Inhibits Calcium-Dependent Neurotransmitter Release. Neuron 95, 577–590 e575

22. Augustin, I., Rosenmund, C., Sudhof, T. C., and Brose, N. (1999) Munc13-1 is essential for fusion competence of glutamatergic synaptic vesicles. Nature 400, 457–461

23. Feldmann, J., Callebaut, I., Raposo, G., Certain, S., Bacq, D., Dumont, C., Lambert, N., Ouachee-Chardin, M., Chedeville, G., Tamary, H., Minard-Colin, V., Vilmer, E., Blanche, S., Le Deist, F., Fischer, A., and de Saint Basile, G. (2003) Munc13-4 is essential for cytolytic granules fusion and is mutated in a form of familial hemophagocytic lymphohistiocytosis (FHL3). Cell 115, 461–473

24. Koch, H., Hofmann, K., and Brose, N. (2000) Definition of Munc13-homology-domains and characterization of a novel ubiquitously expressed Munc13 isoform. Biochem J 349, 247–253

25. Boswell, K. L., James, D. J., Esquibel, J. M., Bruinsma, S., Shirakawa, R., Horiuchi, H., and Martin, T. F. (2012) Munc13-4 reconstitutes calcium-dependent SNARE-mediated membrane fusion. J Cell Biol 197, 301–312

26. Woo, S. S., James, D. J., and Martin, T. F. (2017) Munc13-4 functions as a Ca2+ sensor for homotypic secretory granule fusion to generate endosomal exocytic vacuoles. Mol Biol Cell 28, 792–808

27. Bulfone-Paus, S., Nilsson, G., Draber, P., Blank, U., and Levi-Schaffer, F. (2017) Positive and Negative Signals in Mast Cell Activation. Trends Immunol

28. Theoharides, T. C., Kempuraj, D., Tagen, M., Conti, P., and Kalogeromitros, D. (2007) Differential release of mast cell mediators and the pathogenesis of inflammation. Immunol Rev 217, 65–78

29. McNeil, H. P., Adachi, R., and Stevens, R. L. (2007) Mast cell-restricted tryptases: structure and function in inflammation and pathogen defense. J Biol Chem 282, 20785–20789

30. Reid, G., Wielinga, P., Zelcer, N., van der Heijden, I., Kuil, A., de Haas, M., Wijnholds, J., and Borst, P. (2003) The human multidrug resistance protein MRP4 functions as a prostaglandin efflux transporter and is inhibited by nonsteroidal antiinflammatory drugs. Proc Natl Acad Sci U S A 100, 9244–9249

31. Kochel, T. J., and Fulton, A. M. (2015) Multiple drug resistance-associated protein 4 (MRP4), prostaglandin transporter (PGT), and 15-hydroxyprostaglandin dehydrogenase (15-PGDH) as determinants of PGE2 levels in cancer. Prostaglandins & other lipid mediators 116–117, 99–103

32. Varoqueaux, F., Sigler, A., Rhee, J. S., Brose, N., Enk, C., Reim, K., and Rosenmund, C. (2002) Total arrest of spontaneous and evoked synaptic transmission but normal synaptogenesis in the absence of Munc13-mediated vesicle priming. Proc Natl Acad Sci U S A 99, 9037–9042

33. Scholten, J., Hartmann, K., Gerbaulet, A., Krieg, T., Muller, W., Testa, G., and Roers, A. (2008) Mast cell-specific Cre/loxP-mediated recombination in vivo. Transgenic Res 17, 307–315

34. Schwenk, F., Baron, U., and Rajewsky, K. (1995) A cre-transgenic mouse strain for the ubiquitous deletion of loxP-flanked gene segments including deletion in germ cells. Nucleic Acids Res 23, 5080–5081

35. Makabe-Kobayashi, Y., Hori, Y., Adachi, T., Ishigaki-Suzuki, S., Kikuchi, Y., Kagaya, Y., Shirato, K., Nagy, A., Ujike, A., Takai, T., Watanabe, T., and Ohtsu, H. (2002) The control effect of histamine on body temperature and respiratory function in IgE-dependent systemic anaphylaxis. J Allergy Clin Immunol 110, 298–303

36. Hogan, A. D., and Schwartz, L. B. (1997) Markers of mast cell degranulation. Methods 13, 43–52

37. Fawcett, D. W. (1954) Cytological and pharmacological observations on the release of histamine by mast cells. J Exp Med 100, 217–224

38. Melicoff, E., Sansores-Garcia, L., Gomez, A., Moreira, D. C., Datta, P., Thakur, P., Petrova, Y., Siddiqi, T., Murthy, J. N., Dickey, B. F., Heidelberger, R., and Adachi, R. (2009) Synaptotagmin-2 controls regulated exocytosis but not other secretory responses of mast cells. J Biol Chem 284, 19445–19451

39. Gimborn, K., Lessmann, E., Kuppig, S., Krystal, G., and Huber, M. (2005) SHIP down-regulates FcepsilonR1-induced degranulation at supraoptimal IgE or antigen levels. J Immunol 174, 507–516

40. Kulka, M., Sheen, C. H., Tancowny, B. P., Grammer, L. C., and Schleimer, R. P. (2008) Neuropeptides activate human mast cell degranulation and chemokine production. Immunology 123, 398–410

41. Fernandez, J. M., Neher, E., and Gomperts, B. D. (1984) Capacitance measurements reveal stepwise fusion events in degranulating mast cells. Nature 312, 453–455

42. Alvarez de Toledo, G., and Fernandez, J. M. (1990) Compound versus multigranular exocytosis in peritoneal mast cells. J Gen Physiol 95, 397–409

43. Alvarez de Toledo, G., and Fernandez, J. M. (1990) Patch-clamp measurements reveal multimodal distribution of granule sizes in rat mast cells. J Cell Biol 110, 1033–1039

44. Oberhauser, A. F., and Fernandez, J. M. (1996) A fusion pore phenotype in mast cells of the ruby-eye mouse. Proc Natl Acad Sci U S A 93, 14349–14354

45. Finkelstein, A., Zimmerberg, J., and Cohen, F. S. (1986) Osmotic swelling of vesicles: its role in the fusion of vesicles with planar phospholipid bilayer membranes and its possible role in exocytosis. Annu Rev Physiol 48, 163–174

46. Kasai, H., Takahashi, N., and Tokumaru, H. (2012) Distinct initial SNARE configurations underlying the diversity of exocytosis. Physiol Rev 92, 1915–1964

47. Stevens, R. L., and Adachi, R. (2007) Protease-proteoglycan complexes of mouse and human mast cells and importance of their beta-tryptase-heparin complexes in inflammation and innate immunity. Immunol Rev 217, 155–167

48. Wernersson, S., and Pejler, G. (2014) Mast cell secretory granules: armed for battle. Nat Rev Immunol 14, 478–494

49. Thakurdas, S. M., Melicoff, E., Sansores-Garcia, L., Moreira, D. C., Petrova, Y., Stevens, R. L., and Adachi, R. (2007) The mast cell-restricted tryptase mMCP-6 has a critical immunoprotective role in bacterial infections. J Biol Chem 282, 20809–20815

50. Shin, K., Nigrovic, P. A., Crish, J., Boilard, E., McNeil, H. P., Larabee, K. S., Adachi, R., Gurish, M. F., Gobezie, R., Stevens, R. L., and Lee, D. M. (2009) Mast cells contribute to autoimmune inflammatory arthritis via their tryptase/heparin complexes. J Immunol 182, 647–656

51. Hamilton, M. J., Sinnamon, M. J., Lyng, G. D., Glickman, J. N., Wang, X., Xing, W., Krilis, S. A., Blumberg, R. S., Adachi, R., Lee, D. M., and Stevens, R. L. (2011) Essential role for mast cell tryptase in acute experimental colitis. Proc Natl Acad Sci U S A 108, 290–295

52. Singh, R. K., Mizuno, K., Wasmeier, C., Wavre-Shapton, S. T., Recchi, C., Catz, S. D., Futter, C., Tolmachova, T., Hume, A. N., and Seabra, M. C. (2013) Distinct and opposing roles for Rab27a/Mlph/MyoVa and Rab27b/Munc13-4 in mast cell secretion. FEBS J 280, 892–903

53. Higashio, H., Nishimura, N., Ishizaki, H., Miyoshi, J., Orita, S., Sakane, A., and Sasaki, T. (2008) Doc2 alpha and Munc13-4 regulate Ca(2+)-dependent secretory lysosome exocytosis in mast cells. J Immunol 180, 4774–4784

54. Song, Y., Ailenberg, M., and Silverman, M. (1998) Cloning of a novel gene in the human kidney homologous to rat munc13s: its potential role in diabetic nephropathy. Kidney Int 53, 1689–1695

55. Augustin, I., Betz, A., Herrmann, C., Jo, T., and Brose, N. (1999) Differential expression of two novel Munc13 proteins in rat brain. Biochem J 337, 363–371

56. Rosenmund, C., Sigler, A., Augustin, I., Reim, K., Brose, N., and Rhee, J. S. (2002) Differential control of vesicle priming and short-term plasticity by Munc13 isoforms. Neuron 33, 411–424

57. Dudenhoffer-Pfeifer, M., Schirra, C., Pattu, V., Halimani, M., Maier-Peuschel, M., Marshall, M. R., Matti, U., Becherer, U., Dirks, J., Jung, M., Lipp, P., Hoth, M., Sester, M., Krause, E., and Rettig, J. (2013) Different Munc13 isoforms function as priming factors in lytic granule release from murine cytotoxic T lymphocytes. Traffic 14, 798–809

58. Barbosa, M. D., Nguyen, Q. A., Tchernev, V. T., Ashley, J. A., Detter, J. C., Blaydes, S. M., Brandt, S. J., Chotai, D., Hodgman, C., Solari, R. C., Lovett, M., and Kingsmore, S. F. (1996) Identification of the homologous beige and Chediak-Higashi syndrome genes. Nature 382, 262–265

59. Henningsson, F., Hergeth, S., Cortelius, R., Abrink, M., and Pejler, G. (2006) A role for serglycin proteoglycan in granular retention and processing of mast cell secretory granule components. FEBS J 273, 4901–4912

60. Higashio, H., Satoh, Y., and Saino, T. (2016) Mast cell degranulation is negatively regulated by the Munc13-4-binding small-guanosine triphosphatase Rab37. Sci Rep 6, 22539

61. Pivot-Pajot, C., Varoqueaux, F., de Saint Basile, G., and Bourgoin, S. G. (2008) Munc13-4 regulates granule secretion in human neutrophils. J Immunol 180, 6786–6797

62. Kaeser, P. S., and Regehr, W. G. (2017) The readily releasable pool of synaptic vesicles. Curr Opin Neurobiol 43, 63–70

63. Chen, Z., Cooper, B., Kalla, S., Varoqueaux, F., and Young, S. M., Jr. (2013) The Munc13 proteins differentially regulate readily releasable pool dynamics and calcium-dependent recovery at a central synapse. J Neurosci 33, 8336–8351

64. Dernick, G., de Toledo, G. A., and Lindau, M. (2007) The Patch Amperometry Technique: Design of a Method to Study Exocytosis of Single Vesicles. in Electrochemical Methods for Neuroscience (Michael, A. C., and Borland, L. M. eds.), Boca Raton (FL). pp

65. Tiwari, N., Wang, C. C., Brochetta, C., Ke, G., Vita, F., Qi, Z., Rivera, J., Soranzo, M. R., Zabucchi, G., Hong, W., and Blank, U. (2008) VAMP-8 segregates mast cell-preformed mediator exocytosis from cytokine trafficking pathways. Blood 111, 3665–3674

66. Yu, Y., Huang, Z., Mao, Z., Zhang, Y., Jin, M., Chen, W., Zhang, W., Yu, B., Zhang, W., and Alaster Lau, H. Y. (2016) Go is required for the release of IL-8 and TNF-alpha, but not degranulation in human mast cells. Eur J Pharmacol 780, 115–121

67. Gupta, A. A., Leal-Berumen, I., Croitoru, K., and Marshall, J. S. (1996) Rat peritoneal mast cells produce IFN-gamma following IL-12 treatment but not in response to IgE-mediated activation. J Immunol 157, 2123–2128

68. Leal-Berumen, I., Conlon, P., and Marshall, J. S. (1994) IL-6 production by rat peritoneal mast cells is not necessarily preceded by histamine release and can be induced by bacterial lipopolysaccharide. J Immunol 152, 5468–5476

69. Cairns, J. A., and Walls, A. F. (1996) Mast cell tryptase is a mitogen for epithelial cells. Stimulation of IL-8 production and intercellular adhesion molecule-1 expression. J Immunol 156, 275–283

70. Marshall, J. S., Leal-Berumen, I., Nielsen, L., Glibetic, M., and Jordana, M. (1996) Interleukin (IL)-10 inhibits long-term IL-6 production but not preformed mediator release from rat peritoneal mast cells. J Clin Invest 97, 1122–1128

71. Beil, W. J., Login, G. R., Aoki, M., Lunardi, L. O., Morgan, E. S., Galli, S. J., and Dvorak, A. M. (1996) Tumor necrosis factor alpha immunoreactivity of rat peritoneal mast cell granules decreases during early secretion induced by compound 48/80: an ultrastructural immunogold morphometric analysis. Int Arch Allergy Immunol 109, 383–389

72. Olszewski, M. B., Trzaska, D., Knol, E. F., Adamczewska, V., and Dastych, J. (2006) Efficient sorting of TNF-alpha to rodent mast cell granules is dependent on N-linked glycosylation. Eur J Immunol 36, 997–1008

73. Olszewski, M. B., Groot, A. J., Dastych, J., and Knol, E. F. (2007) TNF trafficking to human mast cell granules: mature chain-dependent endocytosis. J Immunol 178, 5701–5709

74. Dwyer, D. F., Barrett, N. A., Austen, K. F., and Immunological Genome Project, C. (2016) Expression profiling of constitutive mast cells reveals a unique identity within the immune system. Nat Immunol 17, 878–887

75. Grimbaldeston, M. A., Chen, C. C., Piliponsky, A. M., Tsai, M., Tam, S. Y., and Galli, S. J. (2005) Mast cell-deficient W-sash c-kit mutant Kit W-sh/W-sh mice as a model for investigating mast cell biology in vivo. Am J Pathol 167, 835–848

76. Kanaoka, Y., and Boyce, J. A. (2004) Cysteinyl leukotrienes and their receptors: cellular distribution and function in immune and inflammatory responses. J Immunol 173, 1503–1510

77. Adachi, R., Krilis, S. A., Nigrovic, P. A., Hamilton, M. J., Chung, K., Thakurdas, S. M., Boyce, J. A., Anderson, P., and Stevens, R. L. (2012) Ras guanine nucleotide-releasing protein-4 (RasGRP4) involvement in experimental arthritis and colitis. J Biol Chem 287, 20047–20055

78. Kim, K., Petrova, Y. M., Scott, B. L., Nigam, R., Agrawal, A., Evans, C. M., Azzegagh, Z., Gomez, A., Rodarte, E. M., Olkkonen, V. M., Bagirzadeh, R., Piccotti, L., Ren, B., Yoon, J. H., McNew, J. A., Adachi, R., Tuvim, M. J., and Dickey, B. F. (2012) Munc18b is an essential gene in mice whose expression is limiting for secretion by airway epithelial and mast cells. Biochem J 446, 383–394

79. Brock, T. A., Dennis, P. A., Griendling, K. K., Diehl, T. S., and Davies, P. F. (1988) GTP gamma S loading of endothelial cells stimulates phospholipase C and uncouples ATP receptors. Am J Physiol 255, C667–673

80. Lindau, M., and Neher, E. (1988) Patch-clamp techniques for time-resolved capacitance measurements in single cells. Pflugers Arch 411, 137–146

81. Gillis, K. D., Mossner, R., and Neher, E. (1996) Protein kinase C enhances exocytosis from chromaffin cells by increasing the size of the readily releasable pool of secretory granules. Neuron 16, 1209–1220

82. Royet, J. P. (1991) Stereology: a method for analyzing images. Prog Neurobiol 37, 433–474

83. Tschanz, S. A., Burri, P. H., and Weibel, E. R. (2011) A simple tool for stereological assessment of digital images: the STEPanizer. J Microsc 243, 47–59

